# Solving the Pervasive Problem of Protocol Non-Compliance in MRI using an Open-Source tool *mrQA*

**DOI:** 10.1101/2023.07.17.548591

**Authors:** Harsh Sinha, Pradeep Reddy Raamana

## Abstract

Pooling data across diverse sources acquired by multisite consortia requires compliance with a predefined reference protocol *i.e*., ensuring different sites and scanners for a given project have used identical or compatible MR physics parameter values. Traditionally, this has been an arduous and manual process due to difficulties in working with the complicated DICOM standard and lack of resources allocated towards protocol compliance. Moreover, issues of protocol compliance is often overlooked for lack of realization that parameter values are routinely improvised/modified locally at various sites. The inconsistencies in acquisition protocols can reduce SNR, statistical power, and in the worst case, may invalidate the results altogether. An open-source tool, mrQA was developed to automatically assess protocol compliance on standard dataset formats such as DICOM and BIDS, and to study the patterns of non-compliance in over 20 open neuroimaging datasets, including the large ABCD study. The results demonstrate that the lack of compliance is rather pervasive. The frequent sources of non-compliance include but are not limited to deviations in Repetition Time, Echo Time, Flip Angle, and Phase Encoding Direction. It was also observed that GE and Philips scanners exhibited higher rates of non-compliance relative to the Siemens scanners in the ABCD dataset. Continuous monitoring for protocol compliance is strongly recommended before any pre/post-processing, ideally right after the acquisition, to avoid the silent propagation of severe/subtle issues. Although, this study focuses on neuroimaging datasets, the proposed tool mrQA can work with any DICOM-based datasets.

## 1 Introduction

Large-scale neuroimaging datasets play an essential role in characterizing brain-behavior relationships. The average sample size of neuroimaging studies has grown tremendously over the past two decades [1, 2]. Open datasets like the Alzheimer’s Disease Neuroimaging Initiative (ADNI) consists of 800 subjects from 50 sites collected over 2-3 years [3], the Human Connectome Project (HCP) [4] contains 1200 subjects, the Adolescent Brain Cognitive Development (ABCD) study [5] includes over 12000 subjects at 21 sites, the Autism Brain Imaging Data Exchange (ABIDE) provides a dataset of 1000 individuals at 16 international sites, and the UK Biobank is following about 500,000 subjects in the UK. These large-scale datasets are acquired over several years, involving multiple sites, with several vendor-specific scanner models.

A typical MR imaging session consists of multiple modalities (including but not limited to anatomical, functional, and diffusion MRI) along with their corresponding field maps, localizers, and the like for each subject. Imaging data from these modalities provides complementary information about the structural and functional organization. The electronic protocol files generated by scanners (*i*.*e*., Exam Card - Philips, Protocol Exchange - GE, or .exar/.edx file - Siemens) include thousands of parameter values for a single session. To use these distinct modalities effectively, it is important to validate the combinations of acquisition protocols *i*.*e*., evaluating the reliability of chosen imaging sequences and ensuring that the imaging data is acquired accurately for each subject across all sites and scanners. Neither is it a recommended scientific practice nor is it practical to “hope” for data integrity by manual compliance checks across numerous parameters, given the ever-increasing size of neuroimaging studies, cross-site evaluations, multiple scanners, and varied environments.

As maintenance of imaging protocols in MRI centers is typically an ad-hoc and error-prone process, it often leads to variations in acquisition parameters across different subjects and sessions. For instance, manually uploading protocol configurations on each scanner impacts consistency. Inconsistencies also arise from software updates and hardware upgrades which alter the default behavior of the scanning interface.

Even subtle deviations in acquisition parameters can potentially affect the reproducibility of MRI-based brain-behavior studies [6]. Prior works have focused on developing post-processing techniques to reduce the impact of deviations on neuroanatomical estimates [7, 8, 9, 10, 11, 12]. Such post-processing techniques often rely on a large sample size per site to estimate site-specific effects. Recent work by George *et al*. [13] used power analysis to demonstrate that using standardized protocols yields over a two-fold decrease in variability for cortical thickness estimates when compared against non-standardized acquisitions. Therefore, adherence to standardized image acquisition protocols at the scanner is essential for ensuring the quality of MRI-based neuroimaging studies [14, 15, 16]. Otherwise, some subject-specific scans might have to be discarded due to a flawed data collection process, thus reducing the sample size and, consequently, the power of statistical analyses [17]. Yet not much effort has been devoted to eliminating these inconsistencies in image acquisition protocol.

Insufficient monitoring can lead to non-compliance in imaging acquisition parameters, including but not limited to flip angle (FA), repetition time (TR), phase encoding direction (PED), pixel bandwidth (PB), and echo time (TE). When the acquisition parameters are not compliant across scans, it can significantly affect the tissue contrast in T1w/T2w images [18, 19]. In EPI, co-registration with its structural counterpart becomes difficult if EPI is non-compliant with the field map [20, 21]. In DTI, the images acquired with different polarities of PED cannot be used synonymously as they differ in fractional anisotropy estimates [22]. Inconsistencies in image acquisition parameters may implicitly bias the texture in brain images, confounding brain-behavior prediction or phenotypes from brain images [18]. Thus, any analysis conducted without eliminating sources of error in acquisition parameters may reduce statistical power and, in the worst case, may invalidate results altogether, hindering widespread clinical adoption of the experimental results.

Therefore, we present *mrQA* (and *MRdataset*), a software platform to ensure data integrity in MRI datasets. *mrQA* is designed to aggregate and summarize compliance checks on DICOM images at the MRI scanner itself. Automating the compliance check process, *mrQA* can help reduce the risk of errors and omissions in handling and use of DICOM images. DICOM images have an inherent complex structure, and relying on manual interpretation of DICOM fields is prone to error. For instance, left-right flips are not easy to spot visually. However, the ambiguity can be resolved through an automated software that systematically confirms that the DICOM horizontal flip attribute is same as provided in the reference protocol. The software should seamlessly conduct the verification for each scan, removing the necessity for repetitive manual validation [23]. Such subtle errors can have serious consequences, especially for brain surgery. Prior works [24] have proposed validation of acquisition parameters for BIDS datasets. Their work is focused on the execution of BIDS-apps by identifying variations in acquisition parameters. In contrast, *mrQA* focuses on enunciating variation in acquisition parameters for DICOM images. Even though the DICOM format suffers from storage overhead, with complex specifications, DICOM contains complete acquisition metadata with standardized tags. Therefore, it has been the established output format for medical images. In contrast, NIfTI has limited scope for adding important acquisition parameters in the header. The NIfTI format relies on JSON sidecars for storing important acquisition parameters. *mrQA* can discover variations in acquisition parameters in the rawest data format available, *i*.*e*., DICOM format.

*mrQA* has been developed primarily for DICOM-based datasets, but it also expands its functionality to NIfTI-based BIDS datasets. It is important to note that reformatting/validation of BIDS datasets typically occurs years after the data acquisition process has been completed. When non-compliant scans are discovered at a later stage, researchers may have to exclude such subjects/sessions to maintain the reliability of their findings. Therefore, it is important to embrace a mindset of proactive quality assurance *i*.*e*., validating the acquired data as soon as possible to prevent any inconsistencies in acquisition.

An ideal approach is to perform a *real-time* assessment of protocol compliance, which refers to pre-scanning verification of acquisition parameters for compliance during the imaging session, so that scans are not acquired with non-compliant parameters to start with. It might be possible that the default acquisition parameters in the scanning interface are inconsistent with the recommended protocol and real-time checks can help avoid any non-compliance before completing the scan. Although, achieving real-time compliance evaluation is our long-term goal, it is a complex endeavor due to the challenges posed by its logistics and the scanner interfaces. Hence, we focus on evaluating compliance after data-acquisition as a first crucial step to provide a critical perspective on the wide diversity of acquisition parameters in open neuroimaging datasets. It is important to note that this exploration is not about finger-pointing for mistakes. Rather, the motivation is to identify common issues of non-compliance and working collaboratively to address them. Towards this end, we assess protocol compliance, or lack thereof, in the The Adolescent Brain Cognitive Development (ABCD) Study dataset [25], over 20 datasets on OpenNeuro [26] and public DICOM datasets on The Cancer Imaging Archive (TCIA) [27].

## 2 Methods

### 2.1 Overview of *mrQA*

The evaluation of protocol compliance is depicted in two stages as shown in Figure 1. First, we parse the input dataset to create a data structure that stores the acquisition parameters of all the modalities, subjects, and sessions as shown in Figure 2 using *MRdataset* (see Appendix A). Then, the acquisition parameters are aggregated and summarized for generating a protocol compliance report (via *mrQA*). An example script for generating compliance reports is provided in Listing 1. Table 3 provides an example of a compliance report generated for a toy dataset.

**Figure 1:**
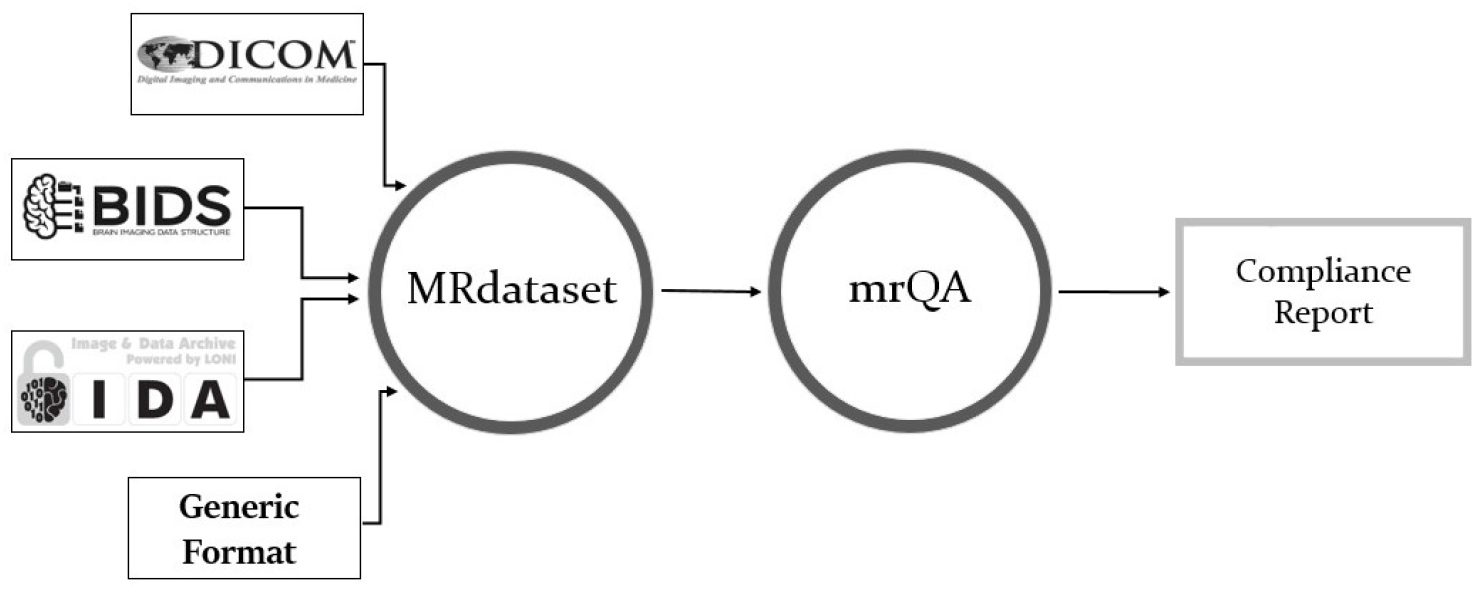
*MRdataset* offers a unified interface to parse & traverse different dataset formats and access acquisition information and metadata e.g. various modalities, subjects, and sessions. This interface is used for generating protocol compliance reports via *mrQA*.

**Figure 2:**
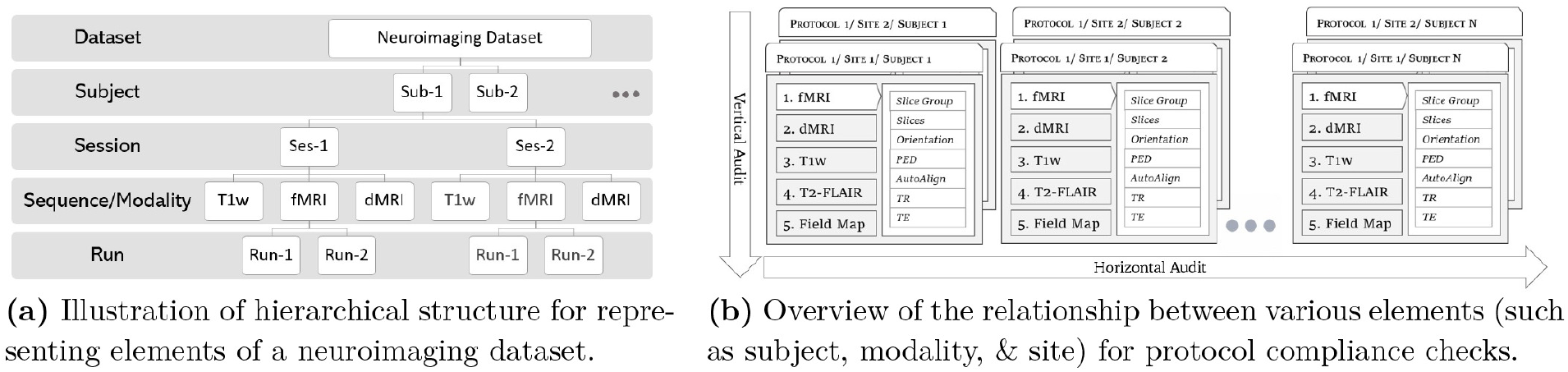
*MRdataset* parses the acquisition parameters for all modalities, subjects, sessions, and runs directly from DICOM headers. Neither does it depend on filename hierarchy nor it expects a particular file organization on disk to accommodate varied configurations in MRI datasets. Then, the parameter values are aggregated to assess protocol compliance for a neuroimaging dataset. We define a horizontal audit to be across all subjects in a given modality (compliant w.r.t a predefined protocol), whereas vertical audit checks if a single subject is compliant across all the acquired modalities.

There can be two types of compliance evaluations - a *horizontal audit* and a *vertical audit*. A *horizontal audit* is focused on assessing parameters for each modality w.r.t. a reference protocol across all subjects in a dataset. A reference protocol is a pre-defined value for each of the acquisition parameters. In a *horizontal audit*, a run is said to be *compliant* if the acquisition parameters for the run are same as the reference protocol. As shown in Figure 2, a subject may have one or more sessions for each modality (e.g. T1w) and each session has multiple runs. A subject is said to be *compliant* for a given modality if all the sessions for the subject are *compliant* with the reference protocol. Therefore, a subject can be *compliant* for one modality (say T1w), but it might be *non-compliant* for another modality (say T2w). A subject is tagged as *non-compliant* even if a single run is found to be non-compliant. A modality is said to be *compliant* if all the subjects in this modality are *compliant* for all sessions. This means there might be some datasets where none of the subjects are compliant.

A *horizontal audit* is essential to ensure the acquisitions across sessions were performed correctly. However, a *horizontal audit* does not address the interaction between multiple modalities within a given session. In contrast, a *vertical audit* checks for compliance issues across all the modalities for each subject within an imaging session. For example, given a subject, all field maps must be set up with the same field-of-view, number of slices, slice thickness, and angulation as the EPI [20]. Similarly, shimming method is specific to a subject [28]. We encourage use of high-order shimming that is consistent across all the subjects in the dataset, especially for spectroscopic experiments [29]. However, minor deviations in shimming across subjects may not warrant the exclusion of a scan. In addition, vertical audits are helpful in revealing specific scans which are found to be non-compliant across multiple modalities. For instance, a vertical audit can spot navigator slices that might have been erroneously uploaded along with a scan for a subject. We recommend that both *horizontal audit* and *vertical audit* must be enforced to eliminate subtle errors in acquisition protocols.

Further, we advocate a two-pronged approach for checking compliance against a reference protocol. The first is *pre-acquisition* compliance, where the parameters will be checked for compliance against a reference protocol before a scan is performed. And the second step is *post-acquisition* compliance, where the parameters are checked after complete data acquisition, validating the *acquired* dataset for compliance. Ideally, both of these two prongs should be performed to maximize data integrity and to minimize loss *i*.*e*., carrying out pre-acquisition compliance checks at initial setup to prevent bad acquisitions in the first place and validating the acquired images with post-acquisition compliance checks to remove any accidental or unknown sources of non-compliance.

In addition, *mrQA* is also being used for continuous monitoring of DICOM datasets in MR labs. *mrQA* can be set up as a cron job to generate reports at regular (daily/weekly) intervals. Meanwhile, if new sessions are acquired, *mrQA* reads the new DICOM files added since the previous run and generates updated compliance reports for the study. The automatic reporting feature is especially useful to notify researchers about any non-compliance in a timely manner so that corrective action can be taken promptly. An example script is provided in Listing 2.

**Listing 1:**
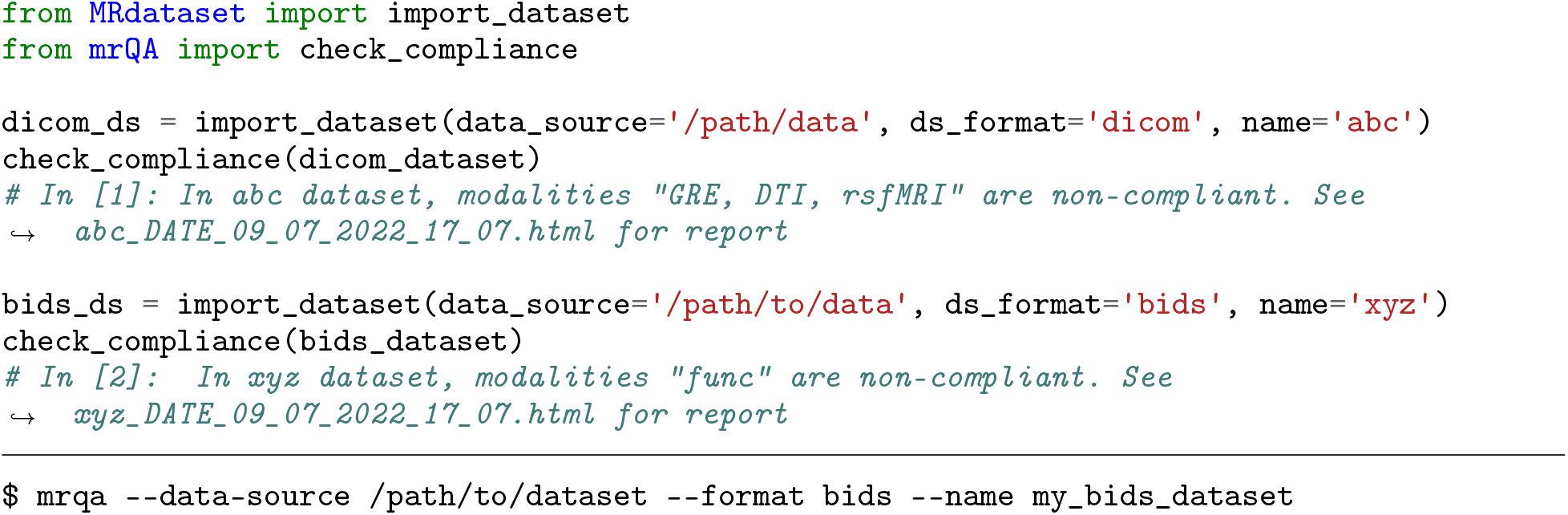
An example using *mrQA* (above: Python API, below: CLI) for generating a compliance report.

### 2.2 Experimental Setup

In this work, we focus on the horizontal audit via post-acquisition compliance to assess neuroimaging datasets for compliance. Assuming that acquisition for most subjects in a given study follows a predefined recommended protocol, *mrQA* infers the most frequent values for each parameter within a modality to construct the *reference protocol*. Then for each subject in the modality, *mrQA* compares the parameter values of each run with the *reference protocol* to determine whether the subject is non-compliant. Finally, each modality is indicated with scores of non-compliance and compliance percentage as shown in Equation 1.

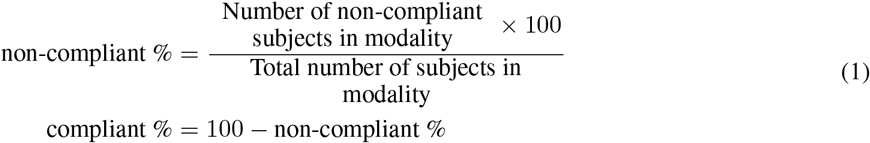

By default, *mrQA* checks for absolute equivalence of parameter values. Although, absolute equivalence is preferred to minimize incongruities, minor differences in decimal values may not necessarily be a part of inclusion/exclusion criteria for a subject. Therefore, we analyze the non-compliance percentage by increasing the tolerance level *i*.*e*., increasing the acceptable range of variation in parameter values against the reference value as shown in Equation 2.

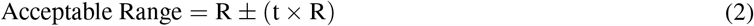

where *R* denotes the parameter value in the reference protocol, and *t* denotes the tolerance level. In this work, we adjust the tolerance level *t* between 0.01 to 0.05. Note that changing the tolerance level will not necessarily decrease the non-compliance rate if the deviations are significant, or if parameters are categorical (*e*.*g*., PED).

We focused on evaluating public datasets as they often serve as a benchmark for neuroimaging analyses. Using *mrQA*, we evaluated three distinct collections of neuroimaging datasets for protocol compliance. First, we evaluated DICOM images from the ABCD Dataset [25] as it provides a unique opportunity to test on a large and diverse sample of over 11,000 subjects (Table 1). Secondly, we utilized 20 large BIDS datasets publicly available on OpenNeuro (Table 2). The datasets were chosen based on their size and availability of JSON sidecar files. Finally, we analyzed DICOM datasets available on The Cancer Imaging Archive (TCIA) (see Appendix C).

**Table 1:** The table summarizes the compliance report for DICOM images from ABCD-baseline FastTrack Active series. For each of the modality (i.e. T1w, T2w, DTI, rsfMRI and field maps), the table shows the vendor, the percentage of non-compliant & compliant subjects, and the parameters which were found to be non-compliant i.e. Repetition Time (TR), Echo Time (TE), Flip Angle (FA) and Pixel Bandwidth (PB). Some minor cases were observed in Phase Encoding Direction (PED), Phase Encoding Steps (PES), Echo Train Length (ETL), and Shim. In contrast to scans acquired with Philips and GE, images scanned with Siemens exhibit minimal non-compliance across all the modalities. Ensuring compliance in acquisition parameters manually is non-trivial for large-scale multi-site datasets such as ABCD. Automated tools like mrQA can help researchers achieve protocol compliance in a practical manner.

| Modality | Vendor | #Non-compliant Subjects |  | Total Subjects | Parameters | #Compliant Subjects |  |
| --- | --- | --- | --- | --- | --- | --- | --- |
| **T1w** | GE | 59 | 2.00 % | 2941 | TE, TR, PB | 2882 | 97.99 % |
|  | Philips | 980 | 64.43 % | 1521 | TE, ETL, PED, PES, PB, TR | 541 | 35.56 % |
|  | Siemens | 2883 | 39.96 %<sup>a</sup> | 7214 | Shim, TE | 4331 | 60.03 % |
| **T2w** | GE | 907 | 32.19 % | 2817 | TE, TR, PB | 1910 | 67.80 % |
|  | Philips | 916 | 62.82 % | 1458 | TE, PB | 542 | 37.17 % |
|  | Siemens | 6 | 0.08 % | 7030 | PED, Shim | 7024 | 99.91 % |
| **Diffusion Fmap** | GE | 620 | 22.26 % | 2785 | FA, PB | 2165 | 77.73 % |
| **Diffusion Fmap**<br>**A≫P** | Philips | 3 | 0.20 % | 1441 | PB | 1438 | 99.79 % |
|  | Siemens | 128 | 1.81 % | 7057 | PED, TR, Shim | 6929 | 98.18 % |
| **Diffusion Fmap**<br>**P≫A** | Philips | 4 | 0.27 % | 1439 | PB | 1435 | 99.72 % |
|  | Siemens | 233 | 3.30 % | 7053 | PED, TR, Shim | 6820 | 96.69 % |
| **fMRI Fmap** | GE | 0 | 0.00 % | 2862 |  | 2862 | 100.00 % |
| **fMRI Fmap**<br>**A≫P** | Philips | 873 | 58.70 % | 1487 | FA, PB | 614 | 41.29 % |
|  | Siemens | 1 | 0.01 % | 7200 | Shim | 7199 | 99.98 % |
| **fMRI Fmap**<br>**P≫A** | Philips | 875 | 58.76 % | 1489 | FA, PB | 614 | 41.23 % |
|  | Siemens | 0 | 0.00 % | 7202 |  | 7202 | 100.00 % |
| **DTI** | GE | 581 | 21.32 % | 2725 | FA, PB | 2144 | 78.67 % |
|  | Philips | 11 | 0.81 % | 1343 | PB | 1332 | 99.18 % |
|  | Siemens | 14 | 0.23 % | 5873 | PED | 5859 | 99.76 % |
| **resting-state**<br>**fMRI** | GE | 674 | 27.16 % | 2481 | PB | 1807 | 72.83 % |
|  | Philips | 787 | 63.98 % | 1230 | PB | 443 | 36.01 % |
|  | Siemens | 2372 | 40.34 % | 5880 | iPAT | 3508 | 59.65 % |

<sup>a</sup> There are minor deviations in TE (ms) within the range (2.88, 2.9)

**Table 2:** The table summarizes compliance report for some OpenNeuro datasets that exhibit deviations in acquisition protocol. For each of these datasets, the table shows the modality, the associated suffix for various tasks/acquisition, the percentage of non-compliant & compliant subjects for each modality, and the parameters which were found to be non-compliant i.e. Repetition Time (TR), Echo Time (TE), and Flip Angle (FA). Some minor cases were observed in Phase Encoding Direction (PED), Phase Encoding Steps (PES), Sequence Variant and Pixel Bandwidth (PB). Thus, mrQA provides the ability to automatically discover scanner-related variance in MR datasets. Automatic compliance checks are especially important for large datasets which exhibit non-compliance rate below 1% because manual/ad-hoc checks are ineffective at detecting these subtle issues.

| Dataset | Modality | Differentiating Entities* | Vendor | #Non-Compliant Subjects |  | Total Subjects | Parameters | #Compliant Subjects |  |
| --- | --- | --- | --- | --- | --- | --- | --- | --- | --- |
| ds000201 | dwi |  | GE | 21 | 27.63% | 76 | PB | 55 | 72.36% |
|  | fmap |  |  | 24 | 28.23% | 85 | PED, PB, TR | 61 | 71.76% |
| ds003826 | anat | t1w | Siemens | 2 | 1.47% | 136 | PES, SV | 134 | 98.52% |
| ds004215 | anat | acq-cube_t2w | GE | 39 | 25.49% | 153 | TE, TR | 114 | 74.50% |
|  | dwi | dir-unflipped |  | 2 | 1.39% | 143 | PB | 141 | 98.60% |
|  | fmap | acq-bold |  | 1 | 1.69% | 59 | PED | 58 | 98.30% |
|  | fmap | acq-dwi |  | 2 | 3.03% | 66 | PED, PB | 64 | 96.96% |
|  | func | task-rest_dir-forward |  | 6 | 4.61% | 130 | FA, PED | 124 | 95.38% |
|  | func | task-rest_dir-reverse |  | 40 | 31.00% | 129 | FA, PED | 89 | 68.99% |
|  | perf | asl |  | 1 | 0.70% | 142 | TR | 141 | 99.29% |
| ds000030 | anat | t1w | Siemens | 92 | 34.71%<sup>†</sup> | 265 | PES, PB | 173 | 65.28% |
|  | dwi |  |  | 112 | 42.74% | 262 | PB, TR | 150 | 57.25% |
|  | func | task-bart |  | 7 | 2.66% | 263 | PED | 256 | 97.33% |
|  | func | task-bht |  | 8 | 3.10% | 258 | PED | 250 | 96.89% |
|  | func | task-pamnec |  | 5 | 2.41% | 207 | PED | 202 | 97.58% |
|  | func | task-pamret |  | 6 | 2.88% | 208 | PED | 202 | 97.11% |
|  | func | task-rest |  | 9 | 3.35% | 268 | PED | 259 | 96.64% |
|  | func | task-scap |  | 9 | 3.35% | 268 | PED | 259 | 96.64% |
|  | func | task-stopsignal |  | 9 | 3.38% | 266 | PED | 257 | 96.61% |
|  | func | task-taskswitch |  | 9 | 3.38% | 266 | PED | 257 | 96.61% |
| ds000221 | fmap | acq-GE | Siemens | 1 | 0.31% | 317 | PED | 316 | 99.68% |
|  | fmap | acq-SE |  | 1 | 0.44% | 227 | PED | 226 | 99.55% |
|  | func | task-rest_acq-PA |  | 14 | 7.07% | 198 | TE | 184 | 92.92% |
| ds002345 | func | task-milkway | Siemens | 17 | 32.07%<sup>†</sup> | 53 | TR | 36 | 67.92% |
| ds000228 | func | task-pixar | Siemens | 3 | 1.93% | 155 | FA | 152 | 98.06% |
| ds000258 | func | task-rest | Siemens | 4 | 4.49%<sup>†</sup> | 85 | TR | 85 | 95.50% |
| ds002785 | dwi |  | Philips | 33 | 15.63%<sup>†</sup> | 211 | TR | 178 | 84.36% |
| ds004169 | anat | t1w | Siemens | 27 | 2.24% | 1202 | FA, PES, PB, TR | 1175 | 97.75% |
|  | func | task-nback |  | 3 | 0.25% | 1189 | FA, PED | 1186 | 99.74% |
|  | func | task-rest |  | 2 | 0.19% | 1029 | FA, PED | 1027 | 99.80% |

<sup>†</sup> There are minor differences in parameter values. See discussion.
\* For **BIDS** datasets, entities correspond to an altered acquisition parameter

We analyzed ABCD-baseline scans for 4 modalities, namely T1w, T2w, DTI, resting-state fMRI, and associated field maps (referred to as fmap), as shown in Table 1. We analyze DICOM images from the ABCD FastTrack Active Series as it closely represents the unprocessed dataset with the most-complete information (closest to the scanners). We assume that all the data collected so far has been acquired with a single protocol as published in Table 2 in Casey *et al*. [5] but we are aware that this protocol might have changed slightly over the years for various reasons. As these details are currently not accessible to us during our analysis of the dataset as a whole, we analyzed it as it was shared. If we redo the analyses accounting for such approved intentional changes in the reference protocol, our results are likely to change and we may see different levels of non-compliance. To accommodate such intentional changes, it is best to run *mrQA* on subsets with a single fixed reference protocol for an accurate estimation of non-compliance in the dataset.

OpenNeuro [26] is a data archive dedicated to open neuroscience data sharing based on FAIR principles [30]. Table 2 presents some of the datasets which exhibit non-compliance in acquisition parameters. Due to the absence of standard acquisition metadata in NIfTI files, we rely on associated JSON sidecar files for evaluating protocol compliance on NIfTI-based datasets.

**Listing 2:**
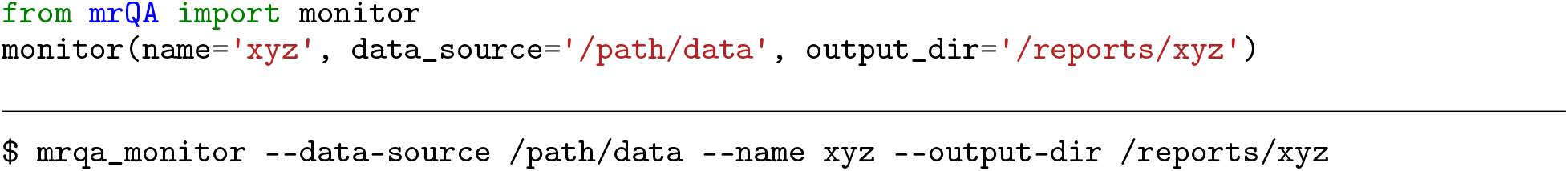
An example of *mrQA* being used for monitoring *xyz* dataset. (above: Python API, below: CLI).

## 3 Results

### 3.1 Evaluation of ABCD dataset

We observed that T1w MRI scans from Philips scanners exhibit non-compliance of 64.43%. As shown in Figure 3, the echo-time (TE) varies in the range (1.4 ms, 3.56 ms) for Philips scanners, even though structural scans are not multi-echo in general. Similarly, T1w images from the GE scanner have minor issues of non-compliance in TE, TR, and PB. T1w scans from Siemens scanners exhibit some minor issues in TE and shim. Although echo time varies for 39.96% of the subjects, there are only minor deviations within the range (2.88 ms, 2.9 ms).

**Figure 3:**
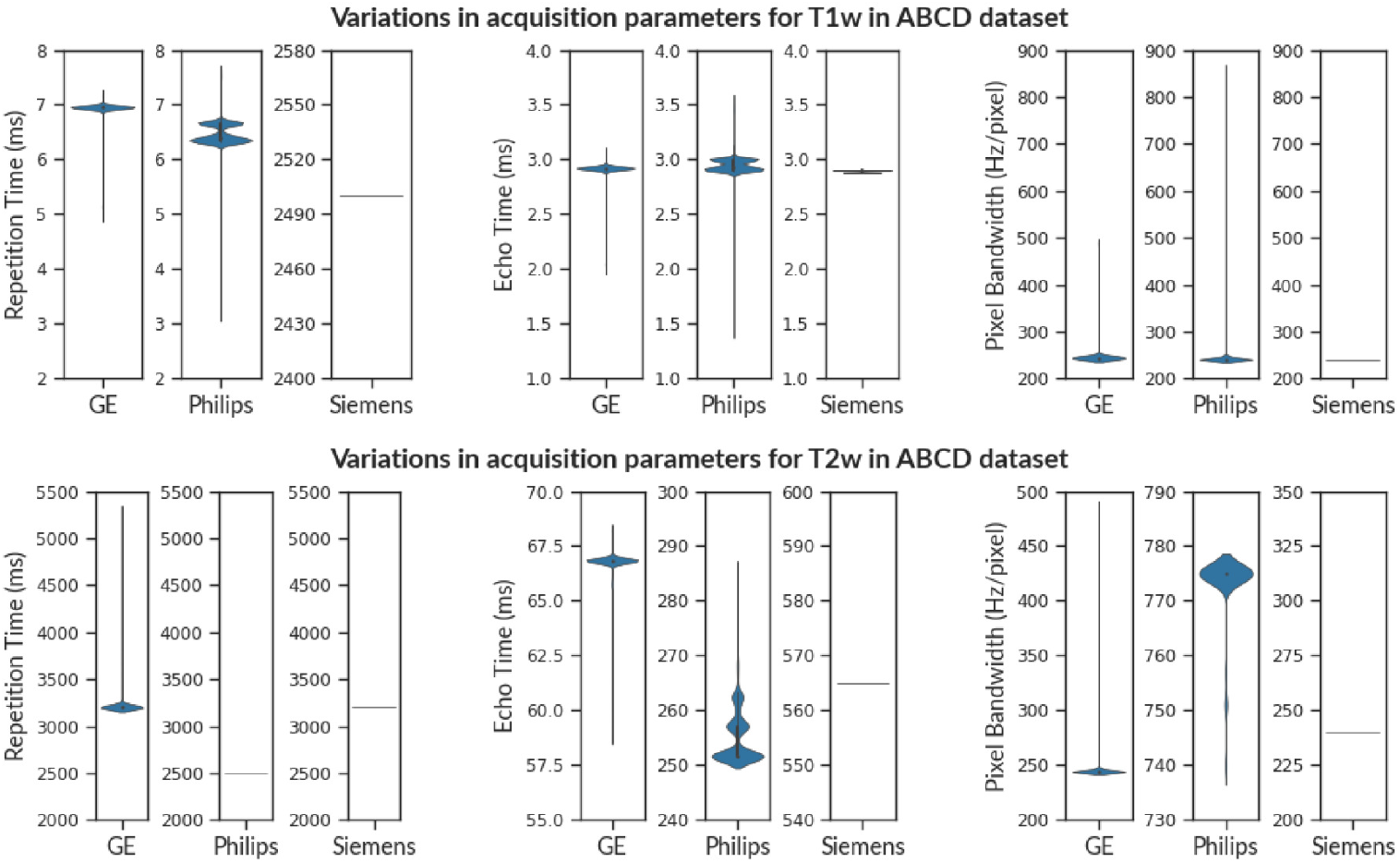
The violin plot shows the variance in Repetition Time (TR), Echo Time (TE), and Pixel Bandwidth (PB) for T1w images (above) and T2w images (below) in the ABCD Dataset. Observe that various vendors have a distinct range of acquisition parameters e.g. Repetition Time (T1w) and Echo Time (T2w). This is because different vendors provide distinct imaging sequences even though the modality might be the same (T1w). Therefore, checking cross-vendor compliance is non-trivial. We observe that scans from Siemens have consistent acquisition parameters in contrast to scans from Philips and GE scanners for both T1w and T2w images.

Similar to T1w images, we observe that 62.82% of subjects are non-compliant for T2w scans from Philips scanners. Figure 3 shows that TE varies in the range (251.49 ms, 285.23 ms) while PB (Hz/pixel) varies between (740, 775). We observe considerable non-compliance (32.19%) in TE, and TR values from GE scanners for T2w images. TE varies in the range (59.1 ms, 68.2 ms) while TR varies in the range (3200 ms, 5297 ms). In contrast, Siemens scanners exhibit minor issues in PED and shim for only 0.08% of subjects.

Table 1 shows the assessment of field maps (fmap) in the ABCD dataset. The subjects are stratified into vendor and PED-specific groups as per information in the DICOM header. Often neuroimaging experiments consist of both A≫P and P≫A scans to reduce susceptibility artifacts[31]. Therefore, the scans will not have a unique PED across all scans. To avoid misinterpretation, compliance checks should be performed within these sub-groups of A≫P and P≫A scans. Note that in the ABCD dataset, Siemens and Philips scanners had distinct field maps each annotated with a PED (AP/PA). However, such annotation was absent in field maps from GE scanners. The field maps intended for Diffusion Images should not be compared to the field maps intended for fMRI images. This information is not automatically captured in DICOM images and should be annotated manually after acquisition.

We observe that both the field maps and Diffusion images from GE scanners have two distinct values of flip angles i.e. 77° and 90°. Even though fMRI field maps from Philips are acquired with flip angle values of 52° and 90°, the resting-state fMRI scans were acquired only with a flip angle of 52°. The report indicates that these subjects don’t comply with a single predefined value for flip angle. We choose to flag this issue, however, whether it is an issue or a study requirement would be best judged by the investigators of the study[32, 33]. In contrast, field maps acquired with Siemens scanner have a flip angle of 90°, and some minor issues in Shim, PED, and TR. We also observed that the Table 2 from Casey *et al*. suggests that parallel imaging was turned off for fMRI sequences acquired with Siemens scanners. But our results show that 60% of subjects were acquired using SENSE.

Figure 4 shows how increasing the relative tolerance level affects the level of non-compliance for T1w, T2w images, and field maps in the ABCD dataset. For T1w and T2w images, the percentage of non-compliance drops close to zero (except for T1w Philips), indicating that the variations lie within 5% tolerance. For T1w from Philips scanners, TE and TR varies beyond the 5% tolerance range of (2.85, 3.15) w.r.t. reference value of 3 ms and (6.33, 6.99) w.r.t. reference value of 6.66 ms, respectively. This results in the non-compliance rate of 22.68% at 5% tolerance level. As the tolerance level is raised from 1% to 5%, the non-compliance rates for diffusion field maps (GE) and fMRI field maps (Philips) show no further decline beyond 22.26% and 35.91%, respectively. This observation can be attributed to large deviations in parameters (such as flip angle and pixel bandwidth) from their reference value, exceeding the 5% tolerance limit.

**Figure 4:**
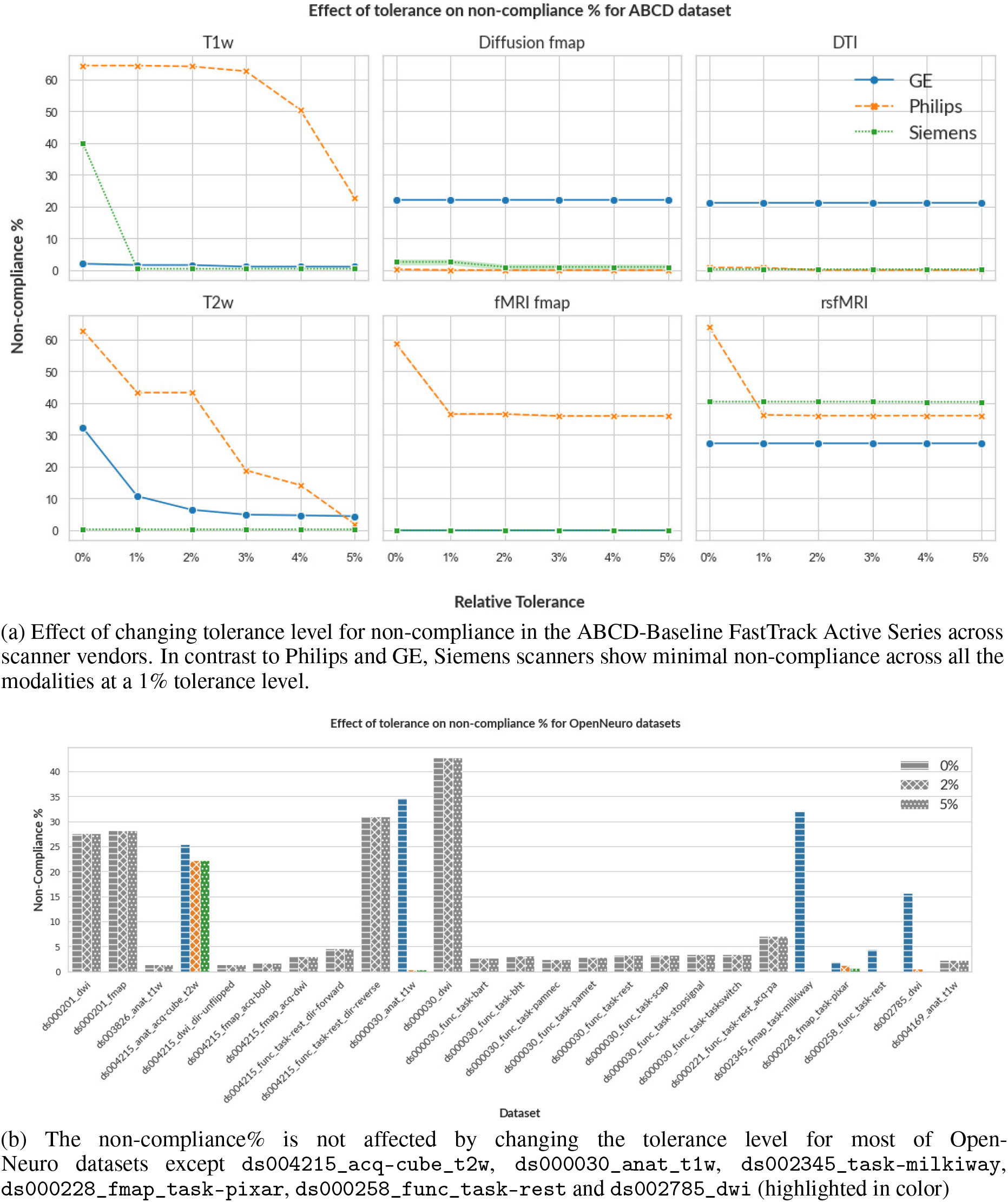
By default, *mrQA* checks for absolute equivalence of parameter values. Given the context of the neuroimaging study, it might be possible to include tolerance in the variation of these acquisition parameters, however, the tolerance level should be best judged by investigators. Note that changing the tolerance level will not affect the non-compliance% if the deviation is too large, or if acquisition parameters are categorical.

### 3.2 Evaluation of OpenNeuro datasets

Table 2 shows evaluation of protocol compliance for OpenNeuro datasets after stratifying modalities by entities such as *task* and *acquisition*. Note that compliance checks should be performed after stratification into coherent clusters as same sequences are often acquired multiple times with varying acquisition parameters for each subject e.g. DTI scans with different PED (A≫P, P≫A) or separate cognitive/behavioral tasks captured with different acquisition protocols in an fMRI study.

We observed that a lot of subjects in datasets such as `ds002843, ds000117, ds000228, ds001242, ds004116, ds003647`, and `ds002345` were missing crucial parameters (such as PED, magnetic field strength, echo train length) from their respective JSON sidecar. We observe that each of the OpenNeuro datasets export a varying set of acquisition parameters because, unlike DICOM tags, JSON sidecar is not standardized. If there is a considerable level of non-compliance, the dataset can be explicitly standardized before it is used for analyses. However, the standardization would have limited validity due to missing acquisition parameters which might impact the reliability of results. Therefore, we recommend that compliance should be checked using DICOM images, which contain complete acquisition metadata with standardized tags.

We also observed that often the same subjects are tagged as non-compliant across several parameters. This can help in identifying consistent patterns in sources of non-compliance. For example, if the same subject is found to be non-compliant for TR and flip angle in T2w FLAIR sequences, this may indicate that the subject was not comfortable inside the scanner, and therefore SAR was adjusted by reducing flip angle and increasing TR value [34]. Therefore, adequate support may be provided to the particular subject during any further scans to ensure compliance. We found this pattern in several datasets such as `ds003826, ds004169, ds000221, ds000030, ds000201, ds004215`, and `ds000258`. Such patterns might also be helpful in identifying particular sites, or scanners that might be the cause of non-compliance.

Figure 6 provides a visual representation of variance in some of the important acquisition parameters such as TE, TR, PB, and PED across a few OpenNeuro datasets. Further, we evaluate the non-compliance of each of these datasets after increasing the tolerance level as shown in Figure 4. We observe that the percentage of non-compliance decreases for 6 datasets only (shown in color), while the percentage of non-compliance for all other datasets is not affected (shown in gray) due to large deviations from the reference beyond the 5% tolerance level or if the non-compliant parameters are categorical (e.g. PED).

## 4 Discussion

We briefly discuss how deviations in acquisition parameters affect images (see Appendix **??** for details). For instance, the flip angle affects the RF signal of the cycle, thereby affecting the signal intensity [35, 36, 37, 38]. Figure 5 shows variation in flip angle in field maps for the ABCD dataset. Similarly, timing parameters (such as TE, and TR) influence tissue specific response in anatomical images and BOLD response in functional images [39, 40, 41].Figure 3 and Figure 6 show the variance of TE and TR for the ABCD dataset and OpenNeuro datasets, respectively. We also observed that datasets (such as `ds004116` and `ds004114`), aggregated images from various field strengths, (4.7 T - 17.15 T) and (3 T - 14.1 T), respectively. In context to texture features, images with varying field strength cannot be used interchangeably [42]. A study may require multiple scans with varying PED to eliminate susceptibility artifacts [43, 44, 31], however it also leads to significant differences in fractional anisotropy estimates [22, 45]. Figure 6 shows a histogram to visualize the apportionment of PED for `ds004215` and `ds000201` datasets.

**Figure 5:**
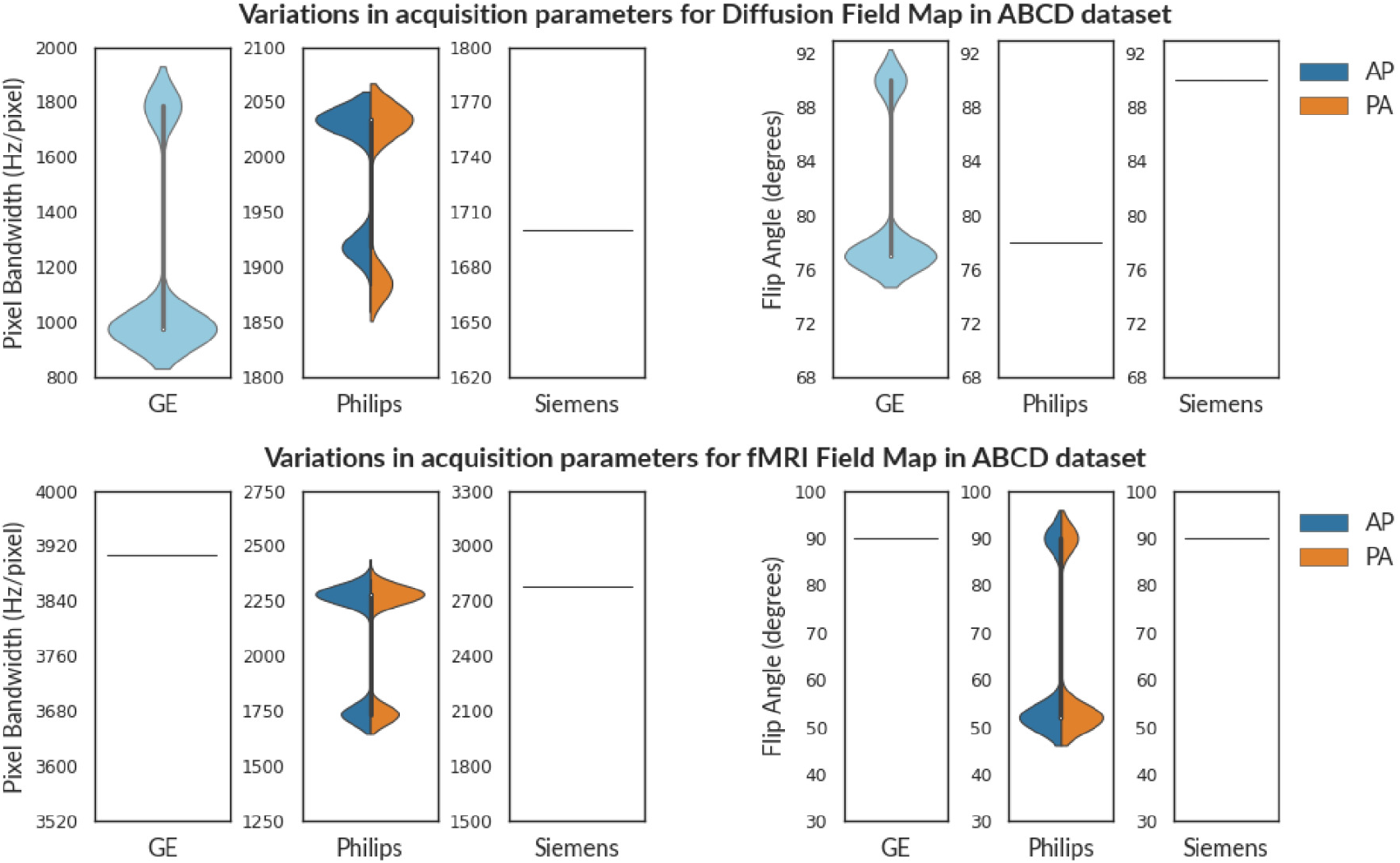
The violin plot shows the variance in Flip Angle(FA) and Pixel Bandwidth (PB) for Diffusion (above) and fMRI (below) field maps in the ABCD Dataset. Siemens and Philips scanners had distinct fieldmaps each annotated with a PED (**AP**/**PA**). However, sequences from GE scanners (denoted by **cyan**) were not annotated in the ABCD dataset. In contrast to scans from Philips and GE scanners, MR scans from Siemens have consistent acquisition parameters across both diffusion and fMRI field maps

**Figure 6:**
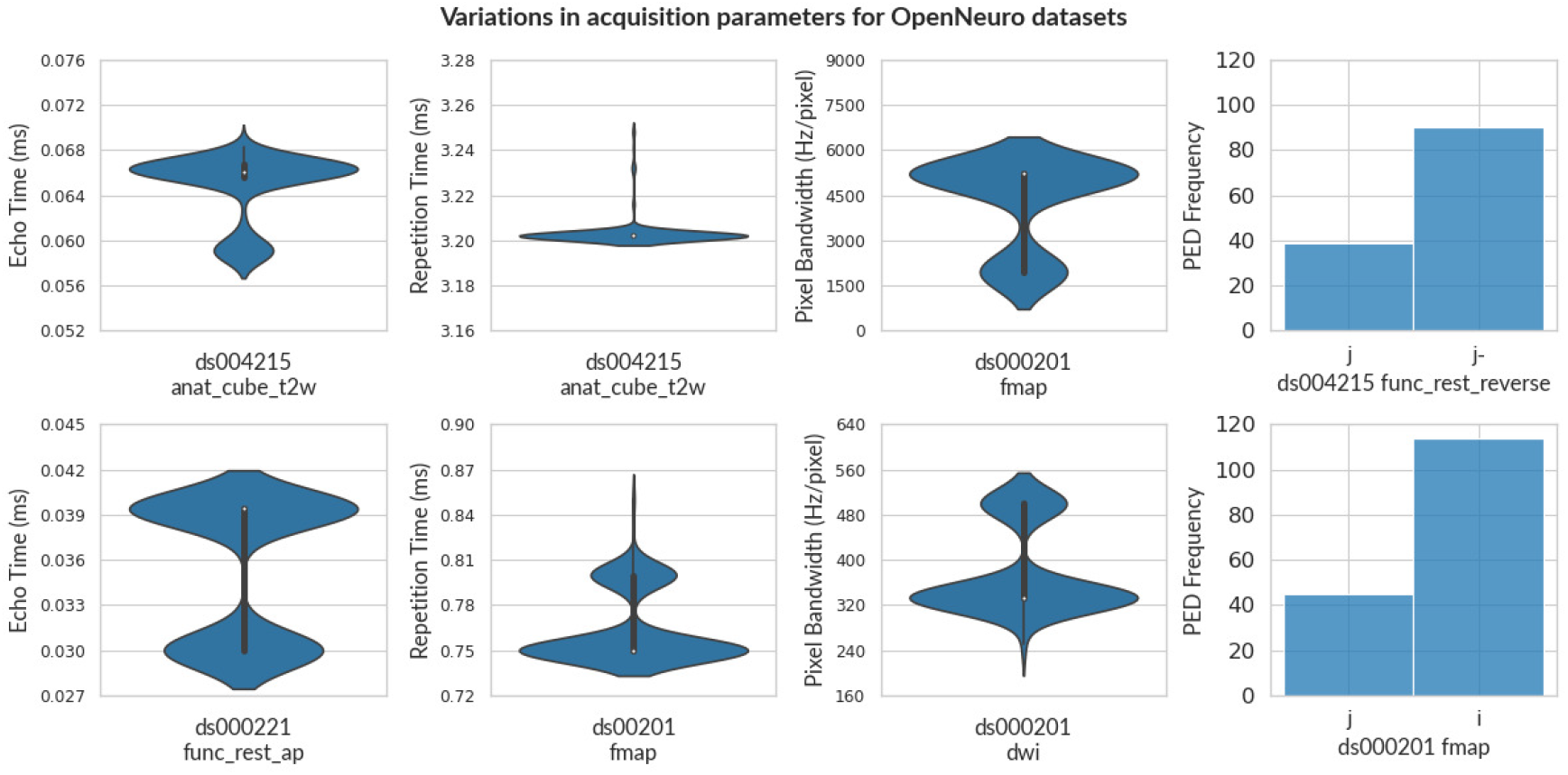
The violin plot shows the variance in Echo Time, Repetition Time, and Pixel Bandwidth for some OpenNeuro Datasets. For violin plots, the width represents the frequency at different levels of each parameter. A histogram chart shows the number of scans for each PED. Note that, even though the entity label specifies PED as *reverse* for `ds004215`, PED is not consistent. This indicates that ensuring compliance is an arduous process, and issues of non-compliance can be overlooked even after careful effort in data acquisition.

Further, we explore the issue of non-conformance in parameters w.r.t. vendors. As compared to Philips and GE scanners, MRI scans acquired with Siemens scanners in the ABCD dataset are observed to be consistent achieving more than 99% compliance over 7000 subjects both in T2w and field maps. We observed that MR scans from Siemens scanners were performed only on Prisma scanners with the same software version (*syngo MR E11*). In contrast, scans for Philips were performed on Achieva dStream and Ingenia models, and the GE scans were executed on MR750 and DV25-26 [5]. Furthermore, both GE and Philips scans had differences in software versions. The difference in hardware and software versions might have been a potential cause of non-compliance in acquisition parameters for Philips and GE scanners. Our findings are consistent with those reported by the ABCD-BIDS Community Collection (ABCC) [46]. They observed a relatively high post-processing quality control failure rate, particularly for images derived from GE and Philips scanners.

Although MRI scanners from different vendors function on the same underlying principles, image sequences can have significant differences in gradient strengths, RF pulse sequences, and timing parameters [47]. In addition, each vendor uses different software and methods to reconstruct the images from *k*-space. Therefore, these sequences are denoted with specific names and abbreviations. For example, Siemens scanners provide an SPACE imaging sequence, while Philips scanners provides VISTA imaging sequence [48]. Both these are 3D TSE sequences and can create T1w images, however significant differences in hardware and software make it non-trivial to compare scans across vendors due to vendor-specific differences. It is better to stratify scans w.r.t. a vendor to avoid any misinterpretations. This problem becomes particularly relevant for multi-site studies where scans are acquired using multiple scanners with potentially differing acquisition protocols. Therefore, the subjects are stratified into different vendor-specific sub-groups in Table 1 as per the information in the DICOM header.

We observed that scanners from various vendors (*e*.*g*., Siemens, GE, Philips) differ in terms of units of measurement and numerical range even for the same parameter. Furthermore, the definition of certain parameters may also vary across vendors based on the particular imaging sequence used. For instance, Field-of-View (FoV) is typically measured in millimeters in Siemens/Philips scanners however GE use centimeters. While analyzing the ABCD dataset, we observed that the TR for Siemens scans was in the range of 2000-4000 ms, but for Philips scans the range was 6-7.5 ms as shown in Figure 3. The precise details of these differences are stored in Exam Card (Philips), Protocol Exchange (GE), or .exar/.edx file (Siemens) generated by corresponding software. Even though much of the information is available in DICOM metadata, inclusion of these electronic protocol files would allow all scanners to have a uniform acquisition protocol loaded into their system without manual intervention, thus eliminating any potential sources of error across various sites and scanners. mrQA is equipped with native support for parsing acquisition protocols in XML files exported from EXAR sources. However, automatic cross-vendor compliance is very difficult due to the lack of standardized open-source tools that can effectively read/write and convert proprietary formats from different vendors.

Finally, we discuss current limitations and future directions for the development of *mrQA. mrQA* extracts certain acquisition parameters such as the shimming method, PAT, and multi-slice mode from Siemens private headers. mrQA skips the private header while reading DICOM images from GE and Philips scanners. Therefore, *mrQA* at its current stage cannot discover non-compliance in parameters present in private headers for GE/Philips scanners.

Note that the DICOM header doesn’t contain any information beyond the specifications of the MR scanner *e*.*g*., variations in duration/intensity of visual stimuli used for measurement of neural responses or reactivity measurements such as CO2 inhalation, acetazolamide infusion [49]. Therefore, checking compliance in the DICOM header may not be sufficient to achieve consistent acquisition.

As of now *mrQA* checks for compliance in a subset of acquisition parameters (in DICOM header) as shown in Table 3. However, there can be other potential sources of non-compliance [50, 41]. Although, this is not necessarily a limitation in itself, as the primary objective of this exploration is to demonstrate the capabilities of automated protocol compliance using *mrQA*(and *MRdataset*). *mrQA* is fully extensible. This means that additional parameters can be readily incorporated into the analysis. As more parameters are integrated, it is indeed likely that the percentage of non-compliance may increase. Nonetheless, the current analysis represents a crucial first step in raising awareness of the prevalence of non-compliance in MR research. By highlighting existing issues and demonstrating the utility of automated compliance assessment tools, we aim to emphasize the imperative need for improved standardization and reporting practices in the field of MR research.

**Table 3:**
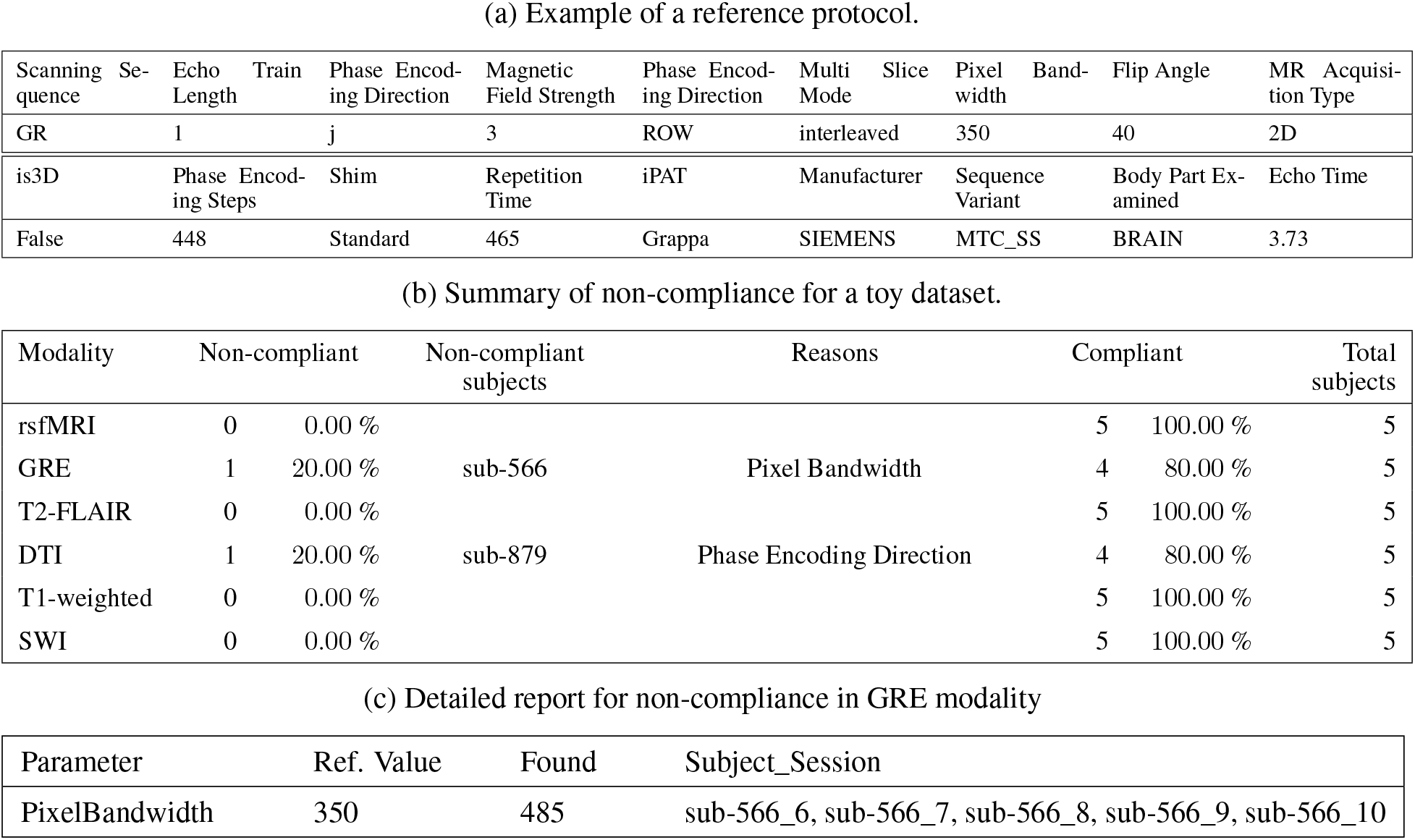
(a) An example reference protocol, and (b,c) compliance report for a toy neuroimaging dataset. Note that the reference protocol can either be pre-defined or inferred by searching for most frequent values for each parameter. The toy dataset has 7 modalities across 5 subjects. Subjects sub-566 and sub-879 are non-compliant w.r.t Pixel Bandwidth and Phase Encoding Direction respectively. For gradient-echo (GRE) modality, subject *sub-566* has a non-compliant pixel bandwidth for sessions 6-10.

## 5 Conclusions

A critical aspect of MR imaging is adherence to the recommended protocol which would enhance the validity and consistency of acquired images. However, we demonstrate the pervasive problem of protocol non-compliance based on analyses of many open datasets from OpenNeuro and the ABCD dataset. Secondly, inconsistencies should be checked promptly so that corrective measures can be taken to minimize differences in acquisition parameters over the entire project timeline. It is non-trivial to maintain protocol compliance in imaging acquisition parameters, especially for large-scale multi-site studies. Monitoring compliance would make us much more familiar with our own data, enabling us to draw meaningful conclusions while considering potential biases, confounds, or anomalies that impact the quality of statistical analysis.

Therefore, we propose an open-source tool, *mrQA* (and *MRdataset*) which can summarize and aggregate acquisition parameters to discover any issues of protocol non-compliance. Apart from generating compliance reports, mrQA can be set up for continuous monitoring of acquired DICOM images on a daily/weekly basis. We believe that it is important to embrace a mindset of proactive quality assurance to weed out any source of inconsistencies at the scanning interface itself rather than waiting for the end-of-analyses to catch confounding. Adopting such an approach before organizing files in a suitable directory structure (e.g. BIDS) will save time and effort.

The long-term goal is to analyze DICOM images in near real-time to identify and fix any issues of non-compliance at the scanner itself. As we move towards even larger datasets, automated imaging QA would be critical for dataset integrity and valid statistical analyses. *mrQA* can help automate this process, as we move towards practical, efficient, and potentially real-time monitoring of protocol compliance.

## Acknowledgments

We would also like to thank Drs. Ashok Panigrahy, Beatriz Luna, Claudiu Schirda, Chan-Hong Moon, Tae Kim, Victor Yushmanov, Andrew Reineberg, Timothy Verstynen and Yaroslav Halchenko for their comments and helpful discussions. We would like to thank Tanupat Boonchalermvichien for his software contributions to the parsing DICOM format. H.S. is supported by the Intelligent Systems Program Fellowship. Pittsburgh Supercomputing Center and the XSEDE initiative provided computational resources.

## Author contributions

Writing - Original Draft Preparation, Figures: H.S.; Writing of the manuscript: H.S. with continuous support from P.R.R in all aspects of writing the manuscript and contributed original ideas. Supervision: P.R.R

## Data Availability Statement

Data sharing is not applicable to this article as no new data were created in this study. Data used in the preparation of this article are publicly available at nda.nih.gov (ABCD), openneuro.org (OpenNeuro) and www.cancerimagingarchive.net (TCIA). The software package is available on the Python Package Manager (PyPI) at https://pypi.org/project/mrQA and its source code is publicly available at https://github.com/Open-Minds-Lab/MRdataset and https://github.com/Open-Minds-Lab/mrQA.

## A Creating a unified interface *(MRdataset)*

Experimental neuroimaging data is hugely diverse and may be structured differently according to study design or clinical protocol, especially the stimulus, behavioral response, and interventions. Thus, any tool must address the fundamental capability of representing and manipulating common dataset formats. *MRdataset* provides a unified interface to simplify the traversal of neuroimaging datasets by adopting modular classes for each dataset format as well as different levels in the hierarchy, such as modalities, subjects, sessions, and runs [51] as shown in Figure 1. *MRdataset* infers information about the hierarchical structure directly from the DICOM headers. *MRdataset* doesn’t rely on filenames or expect a particular idiosyncratic organization of files to process various configurations. It provides a simple modular interface to improve the use and accessibility of neuroimaging datasets.

MRdataset provides a consistent set of methods for data access irrespective of the dataset format (such as BIDS, and DICOM). In the future, we expect to extend the *MRdataset* dataset class to support other data formats, such as LONI IDA, but these were not included in the initial design. In addition to the desired unified interface, *MRdataset* performs basic validation to reject localizers, head scouts, and phantoms. Keeping these considerations in mind, the package is written in Python, and it uses *pydicom* [52] for reading DICOM images. Python is becoming the de facto standard for scientific applications as it provides community support and easy extensibility for the future [53].

## B Example of a compliance report

An HTML report is generated for a toy dataset that presents complete information in a concise manner (as shown in Table 3). The report has two parts - a summary view which gives a brief assessment of protocol compliance for all the modalities. Following the summary, each modality is accompanied by a detailed view. There are two non-compliant subjects, *sub-566* and *sub-879*, for the modalities GRE and DTI, respectively. Subject *sub-566* has non-compliant pixel bandwidth, while subject *sub-879* has an non-compliant PED. For instance, pixel bandwidth in the reference protocol is 350, but subject *sub-566* has a pixel bandwidth of 485 in sessions 6, 7, 8, 9, and 10. The complete compliance report contains reference protocols for each modality and corresponding details about sources of error in acquisition parameters.

## C Evaluation of DICOM datasets on TCIA

Although we focus primarily on neuroimaging datasets, *mrQA* can analyze DICOM-based datasets for other organs also. We also analyze three public DICOM datasets available on The Cancer Imaging Archive (TCIA) [27], namely Rembrandt [54], TCGA-GBM [55] and TCGA-LGG [56]. For Rembrandt, we observe issues of non-compliance in TR. Many scans had missing values for crucial parameters like magnetic field strength, pixel bandwidth, PED for FLAIR, and diffusion images. We observe issues in repetition time, pixel bandwidth, and magnetic field strength for TCGA-LGG and TCGA-GBM datasets. Missing acquisition parameters can not only limit reproducibility, it also limits the ability to perform comparative analysis across studies, which consequently affects the validity and reliability of the research. The compliance reports for TCIA are available at https://github.com/Open-Minds-Lab/mrQA-reports

## D Effect of non-compliance in acquisition parameters

In this section, we discuss the impact on image quality due to variation in specific acquisition parameters.

### D.1 Flip Angle

The flip angle affects the net magnetization relative to the primary magnetic field as it controls the amount of longitudinal magnetization converted to transverse magnetization. Thus, the flip angle affects the RF signal of the cycle, thereby affecting the signal intensity. For instance, large flip angles produce T1 contrast, low flip angles produce PD (proton-density) contrast, and the T1 and PD contrast can cancel each other for intermediate flip angles [35, 36, 37]. Apart from anatomical MRI, the flip angle also affects functional MRI. Large flip angles deteriorate the spatial contrast between cerebrospinal fluid (CSF), gray matter (GM), and white matter (WM), which makes it difficult to align EPI to its structural counterpart [38]. The impact on signal intensity and its significance for pattern discrimination would be governed by specific tissue in context. It is crucial to recognize that flip angles directly influence image contrast.

### D.2 Echo Time & Repetition Time

A correct choice for echo time (TE) and repetition time (TR) is important for structural images. The tissue-specific response can be influenced by acquisition parameters such as TR and TE as different tissues have varying T1 and T2 times. For instance, to generate a valid T1-weighted image, it is important that TR & TE is less than tissue-specific T1 & T2 times, respectively. In contrast, TR & TE should be much greater than tissue-specific T1 time & T2 time, respectively, to generate T2-weighted images. However, if TR is much greater than tissue-specific T1 time but TE is less than tissue-specific T2 time, the result is a proton density-weighted image. Thus, variations in TR and TE can dictate image contrast characteristics [19].

Apart from structural images, choice of TR and TE also affects fMRI images. Longer TE values lead to increased susceptibility artifacts and signal dropout. Shorter TRs provide enhanced BOLD senstivity but may also lead to saturation effects [39, 40, 41].

### D.3 Magnetic Field Strength

Variation in magnetic field strength has a strong influence on texture features (*e*.*g*., co-occurence matrix and gray-level run length matrix). These texture features capture patterns and provide valuable information about visual appearance and structural characteristics of an image. These texture features can significantly impact the performance of predictive models.

Ammari et. al [42] show a comprehensive analysis studying the impact of magnetic field strength on various texture features. The study evaluates 38 texture features of which 15 features in healthy volunteers were sensitive to variations in magnetic field strength. Visually, images from various field strengths may have the same visual diagnostic accuracy [57], but in context to texture features, images with varying magnetic field strength cannot be used interchangeably [42].

### D.4 Phase Encoding Direction

PED plays a crucial role in EPI sequences. A common issue in EPI sequences is their vulnerability to susceptibility artifacts. These artifacts are apparent, especially when the bandwidth is low [43, 44]. To diminish the effect of these distortions, the collection of additional scans with varying PED is very helpful [31]. Although it is important to eliminate these artifacts to improve image quality in DTI sequences, varying the PED is known to affect fractional anisotropy estimates. Kennis et. al [22] show that magnitude of fractional anisotropy magnitude between P≫A and A≫P scans can range from 0.4% to 30% even after correction for subject motion, eddy currents effects, and susceptibility artifacts. It is possible that these differences are arising due to signal intensities from P≫A and A≫P scans that can confound DTI neuroanatomical studies. Similarly, Tudela et. al [45] show that misalignment in PED from the main magnetic field can lead to much more artifacts reflected by lower fractional anisotropy values.

### D.5 Discussion

Prior works have measured the effect of various acquisition protocols on texture analysis [58, 59, 60], to evaluate which features are stable against changes in acquisition protocols. Some parameters such as TR and TE do not affect the shape and size of the image, but they affect uniformity in grayscale intensity [18]. It is evident that different acquisition protocols can affect data distribution, reducing the reliability of the extracted features and consequently increasing the bias of downstream statistical analyses [61]. Thus, special attention must be attributed to image pre-processing before any feature extraction [62].

Harmonization is possible only if we know that the data exhibits variation in imaging acquisition protocol. In cases when sources of non-compliance are not known, data cannot be categorized into clusters, and it would be difficult to perform data harmonization. Thus, *mrQA* can play a pivotal role in establishing data integrity by discovering sources of non-compliance allowing the investigators to perform harmonization, if required.

As we progress towards algorithms that are able to learn features automatically (e.g. deep learning), it is even more important to ensure that the derived image features are stable with respect to variations in acquisition parameters [18]. Without acknowledging these sources of variation in acquisition parameters, the statistical results might be subject to confounding which can obscure or exaggerate the effects of interest [63], leading to misinterpretation of statistical results.

## References

[1] Peter A. Bandettini. Twenty years of functional MRI: The science and the stories. NeuroImage, 62(2):575–588, August 2012. ISSN 10538119. doi:10.1016/j.neuroimage.2012.04.026. URL https://linkinghub.elsevier.com/retrieve/pii/S1053811912004223.

[2] Denes Szucs and John PA. Ioannidis. Sample size evolution in neuroimaging research: An evaluation of highly-cited studies (1990–2012) and of latest practices (2017–2018) in high-impact journals. NeuroImage, 221: 117164, November 2020. ISSN 10538119. doi:10.1016/j.neuroimage.2020.117164. URL https://linkinghub.elsevier.com/retrieve/pii/S1053811920306509.

[3] R. C. Petersen, P. S. Aisen, L. A. Beckett, M. C. Donohue, A. C. Gamst, D. J. Harvey, C. R. Jack, W. J. Jagust, L. M. Shaw, A. W. Toga, J. Q. Trojanowski, and M. W. Weiner. Alzheimer’s Disease Neuroimaging Initiative (ADNI): Clinical characterization. Neurology, 74(3):201–209, January 2010. ISSN 0028-3878, 1526-632X. doi:10.1212/WNL.0b013e3181cb3e25. URL https://www.neurology.org/lookup/doi/10.1212/WNL.0b013e3181cb3e25.

[4] David C. Van Essen, Stephen M. Smith, Deanna M. Barch, Timothy E.J. Behrens, Essa Yacoub, and Kamil Ugurbil. The WU-Minn Human Connectome Project: An overview. NeuroImage, 80:62–79, October 2013. ISSN 10538119. doi:10.1016/j.neuroimage.2013.05.041. URL https://linkinghub.elsevier.com/retrieve/pii/S1053811913005351.

[5] B.J. Casey, Tariq Cannonier, May I. Conley, Alexandra O. Cohen, Deanna M. Barch, Mary M. Heitzeg, Mary E. Soules, Theresa Teslovich, Danielle V. Dellarco, Hugh Garavan, Catherine A. Orr, Tor D. Wager, Marie T. Banich, Nicole K. Speer, Matthew T. Sutherland, Michael C. Riedel, Anthony S. Dick, James M. Bjork, Kathleen M. Thomas, Bader Chaarani, Margie H. Mejia, Donald J. Hagler, M. Daniela Cornejo, Chelsea S. Sicat, Michael P. Harms, Nico U.F. Dosenbach, Monica Rosenberg, Eric Earl, Hauke Bartsch, Richard Watts, Jonathan R. Polimeni, Joshua M. Kuperman, Damien A. Fair, and Anders M. Dale. The Adolescent Brain Cognitive Development (ABCD) study: Imaging acquisition across 21 sites. Developmental Cognitive Neuroscience, 32:43–54, August 2018. ISSN 18789293. doi:10.1016/j.dcn.2018.03.001. URL https://linkinghub.elsevier.com/retrieve/pii/S1878929317301214.

[6] Jorge Jovicich, Silvester Czanner, Xiao Han, David Salat, Andre van der Kouwe, Brian Quinn, Jenni Pacheco, Marilyn Albert, Ronald Killiany, and Deborah Blacker. MRI-derived measurements of human subcortical, ventricular and intracranial brain volumes: Reliability effects of scan sessions, acquisition sequences, data analyses, scanner upgrade, scanner vendors and field strengths. NeuroImage, 46(1):177–192, May 2009. ISSN 10538119. doi:10.1016/j.neuroimage.2009.02.010. URL https://linkinghub.elsevier.com/retrieve/pii/S1053811909001505.

[7] Lee Friedman, Hal Stern, Gregory G. Brown, Daniel H. Mathalon, Jessica Turner, Gary H. Glover, Randy L. Gollub, John Lauriello, Kelvin O. Lim, Tyrone Cannon, Douglas N. Greve, Henry Jeremy Bockholt, Aysenil Belger, Bryon Mueller, Michael J. Doty, Jianchun He, William Wells, Padhraic Smyth, Steve Pieper, Seyoung Kim, Marek Kubicki, Mark Vangel, and Steven G. Potkin. Test-retest and between-site reliability in a multicenter fMRI study. Human Brain Mapping, 29(8):958–972, August 2008. ISSN 10659471. doi:10.1002/hbm.20440. URL https://onlinelibrary.wiley.com/doi/10.1002/hbm.20440.

[8] Sylvain Gouttard, Martin Styner, Marcel Prastawa, Joseph Piven, and Guido Gerig. Assessment of Reliability of Multi-site Neuroimaging Via Traveling Phantom Study. Springer, 5242:263–270, September 2008. doi:10.1007/978-3-540-85990-1_32. URL http://link.springer.com/10.1007/978-3-540-85990-1_32. Series Title: Lecture Notes in Computer Science.

[9] Jorge Jovicich, Silvester Czanner, Douglas Greve, Elizabeth Haley, Andre van der Kouwe, Randy Gollub, David Kennedy, Franz Schmitt, Gregory Brown, James MacFall, Bruce Fischl, and Anders Dale. Reliability in multi-site structural MRI studies: Effects of gradient non-linearity correction on phantom and human data. NeuroImage, 30(2):436–443, April 2006. ISSN 10538119. doi:10.1016/j.neuroimage.2005.09.046. URL https://linkinghub.elsevier.com/retrieve/pii/S1053811905007299.

[10] Heath Pardoe, Gaby S. Pell, David F. Abbott, Anne T. Berg, and Graeme D. Jackson. Multi-site voxel-based morphometry: Methods and a feasibility demonstration with childhood absence epilepsy. NeuroImage, 42(2): 611–616, August 2008. ISSN 10538119. doi:10.1016/j.neuroimage.2008.05.007. URL https://linkinghub.elsevier.com/retrieve/pii/S1053811908006174.

[11] Hugo G. Schnack, Neeltje E.M. van Haren, Hilleke E. Hulshoff Pol, Marco Picchioni, Matthias Weisbrod, Heinrich Sauer, Tyrone Cannon, Matti Huttunen, Robin Murray, and Ren S. Kahn. Reliability of brain volumes from multicenter MRI acquisition: A calibration study. Human Brain Mapping, 22(4):312–320, August 2004. ISSN 1065-9471, 1097-0193. doi:10.1002/hbm.20040. URL https://onlinelibrary.wiley.com/doi/10.1002/hbm.20040.

[12] Jean-Philippe Fortin, Nicholas Cullen, Yvette I. Sheline, Warren D. Taylor, Irem Aselcioglu, Philip A. Cook, Phil Adams, Crystal Cooper, Maurizio Fava, Patrick J. McGrath, Melvin McInnis, Mary L. Phillips, Madhukar H. Trivedi, Myrna M. Weissman, and Russell T. Shinohara. Harmonization of cortical thickness measurements across scanners and sites. NeuroImage, 167:104–120, February 2018. ISSN 10538119. doi:10.1016/j.neuroimage.2017.11.024. URL https://linkinghub.elsevier.com/retrieve/pii/S105381191730931X.

[13] Allan George, Ruben Kuzniecky, Henry Rusinek, Heath R. Pardoe, and for the Human Epilepsy Project Investigators. Standardized Brain MRI Acquisition Protocols Improve Statistical Power in Multicenter Quantitative Morphometry Studies. Journal of Neuroimaging, 30(1):126–133, January 2020. ISSN 1051-2284, 1552-6569. doi:10.1111/jon.12673. URL https://onlinelibrary.wiley.com/doi/10.1111/jon.12673.

[14] Clifford R. Jack, Matt A. Bernstein, Nick C. Fox, Paul Thompson, Gene Alexander, Danielle Harvey, Bret Borowski, Paula J. Britson, Jennifer L. Whitwell, Chadwick Ward, Anders M. Dale, Joel P. Felmlee, Jeffrey L. Gunter, Derek L.G. Hill, Ron Killiany, Norbert Schuff, Sabrina Fox-Bosetti, Chen Lin, Colin Studholme, Charles S. DeCarli, Gunnar Krueger, Heidi A. Ward, Gregory J. Metzger, Katherine T. Scott, Richard Mallozzi, Daniel Blezek, Joshua Levy, Josef P. Debbins, Adam S. Fleisher, Marilyn Albert, Robert Green, George Bartzokis, Gary Glover, John Mugler, Michael W. Weiner, and ADNI Study. The Alzheimer’s disease neuroimaging initiative (ADNI): MRI methods. Journal of Magnetic Resonance Imaging, 27(4):685–691, April 2008. ISSN 10531807, 15222586. doi:10.1002/jmri.21049. URL https://onlinelibrary.wiley.com/doi/10.1002/jmri.21049.

[15] G. Pearlson. Multisite Collaborations and Large Databases in Psychiatric Neuroimaging: Advantages, Problems, and Challenges. Schizophrenia Bulletin, 35(1):1–2, January 2009. ISSN 0586-7614, 1745-1701. doi:10.1093/schbul/sbn166. URL https://academic.oup.com/schizophreniabulletin/article-lookup/doi/10.1093/schbul/sbn166.

[16] Christopher L. Schlett, Thomas Hendel, Jochen Hirsch, Sabine Weckbach, Svenja Caspers, Jeanette Schulz-Menger, Till Ittermann, Florian Von Knobelsdorff-Brenkenhoff, Susanne C. Ladd, Susanne Moebus, Christian Stroszczynski, Beate Fischer, Michael Leitzmann, Christiane Kuhl, Frank Pessler, Dagmar Hartung, Yvonne Kemmling, Holger Hetterich, Katrin Amunts, Matthias Günther, Frank Wacker, Ernst Rummeny, Hans-Ulrich Kauczor, Michael Forsting, Henry Völzke, Norbert Hosten, Maximilian F. Reiser, and Fabian Bamberg. Quantitative, organ-specific interscanner and intrascanner variability for 3 t whole-body magnetic resonance imaging in a multicenter, multivendor study:. Investigative Radiology, 51(4):255–265, Apr 2016. ISSN 0020-9996. doi:10.1097/RLI.0000000000000237.

[17] Katherine S. Button, John P. A. Ioannidis, Claire Mokrysz, Brian A. Nosek, Jonathan Flint, Emma S. J. Robinson, and Marcus R. Munafò. Power failure: why small sample size undermines the reliability of neuroscience. Nature Reviews Neuroscience, 14(5):365–376, May 2013. ISSN 1471-003X, 1471-0048. doi:10.1038/nrn3475. URL http://www.nature.com/articles/nrn3475.

[18] Marius E. Mayerhoefer, Pavol Szomolanyi, Daniel Jirak, Andrzej Materka, and Siegfried Trattnig. Effects of MRI acquisition parameter variations and protocol heterogeneity on the results of texture analysis and pattern discrimination: An application-oriented study: Effects of MRI acquisition parameters on texture analysis. Medical Physics, 36(4):1236–1243, March 2009. ISSN 00942405. doi:10.1118/1.3081408. URL http://doi.wiley.com/10.1118/1.3081408.

[19] Garry E. Gold, Eric Han, Jeff Stainsby, Graham Wright, Jean Brittain, and Christopher Beaulieu. Musculoskeletal MRI at 3.0 T: Relaxation Times and Image Contrast. American Journal of Roentgenology, 183(2):343–351, August 2004. ISSN 0361-803X, 1546-3141. doi:10.2214/ajr.183.2.1830343. URL https://www.ajronline.org/doi/10.2214/ajr.183.2.1830343.

[20] Sijia Wang, Daniel J. Peterson, J. C. Gatenby, Wenbin Li, Thomas J. Grabowski, and Tara M. Madhyastha. Evaluation of field map and nonlinear registration methods for correction of susceptibility artifacts in diffusion mri. Frontiers in Neuroinformatics, 11, 2017. ISSN 1662-5196. URL https://www.frontiersin.org/articles/10.3389/fninf.2017.00017.

[21] Peter Jezzard. Correction of geometric distortion in fMRI data. NeuroImage, 62(2):648–651, August 2012. ISSN 10538119. doi:10.1016/j.neuroimage.2011.09.010. URL https://linkinghub.elsevier.com/retrieve/pii/S105381191101055X.

[22] M. Kennis, S.J.H. van Rooij, R.S. Kahn, E. Geuze, and A. Leemans. Choosing the polarity of the phase-encoding direction in diffusion MRI: Does it matter for group analysis? NeuroImage: Clinical, 11:539–547, 2016. ISSN 22131582. doi:10.1016/j.nicl.2016.03.022. URL https://linkinghub.elsevier.com/retrieve/pii/S2213158216300638.

[23] Daniel R. Glen, Paul A. Taylor, Bradley R. Buchsbaum, Robert W. Cox, and Richard C. Reynolds. Beware (Surprisingly Common) Left-Right Flips in Your MRI Data: An Efficient and Robust Method to Check MRI Dataset Consistency Using AFNI. Frontiers in Neuroinformatics, 14:18, May 2020. ISSN 1662-5196. doi:10.3389/fninf.2020.00018. URL https://www.frontiersin.org/article/10.3389/fninf.2020.00018/full.

[24] Sydney Covitz, Tinashe M. Tapera, Azeez Adebimpe, Aaron F. Alexander-Bloch, Maxwell A. Bertolero, Eric Feczko, Alexandre R. Franco, Raquel E. Gur, Ruben C. Gur, Timothy Hendrickson, Audrey Houghton, Kahini Mehta, Kristin Murtha, Anders J. Perrone, Tim Robert-Fitzgerald, Jenna M. Schabdach, Russell T Shinohara, Jacob W. Vogel, Chenying Zhao, Damien A. Fair, Michael P. Milham, Matthew Cieslak, and Theodore D. Satterthwaite. Curation of bids (cubids): A workflow and software package for streamlining reproducible curation of large bids datasets. NeuroImage, 263:119609, Nov 2022. ISSN 10538119. doi:10.1016/j.neuroimage.2022.119609.

[25] Terry L Jernigan. Adolescent Brain Cognitive Development Study (ABCD), January 2017. URL https://nda.nih.gov/edit_collection.html?id=2573.

[26] Christopher J Markiewicz, Krzysztof J Gorgolewski, Franklin Feingold, Ross Blair, Yaroslav O Halchenko, Eric Miller, Nell Hardcastle, Joe Wexler, Oscar Esteban, Mathias Goncavles, Anita Jwa, and Russell Poldrack. The openneuro resource for sharing of neuroscience data. eLife, 10:e71774, Oct 2021. ISSN 2050-084X. doi:10.7554/eLife.71774.

[27] Kenneth Clark, Bruce Vendt, Kirk Smith, John Freymann, Justin Kirby, Paul Koppel, Stephen Moore, Stanley Phillips, David Maffitt, Michael Pringle, Lawrence Tarbox, and Fred Prior. The Cancer Imaging Archive (TCIA): Maintaining and Operating a Public Information Repository. Journal of Digital Imaging, 26(6):1045– 1057, December 2013. ISSN 0897-1889, 1618-727X. doi:10.1007/s10278-013-9622-7. URL http://link.springer.com/10.1007/s10278-013-9622-7.

[28] Rolf Gruetter. Automatic, localizedin Vivo adjustment of all first-and second-order shim coils. Magnetic Resonance in Medicine, 29(6):804–811, June 1993. ISSN 07403194, 15222594. doi:10.1002/mrm.1910290613. URL https://onlinelibrary.wiley.com/doi/10.1002/mrm.1910290613.

[29] Peter B Barker, Alberto Bizzi, Nicola De Stefano, Doris DM Lin, and Rao Gullapalli. Clinical MR spectroscopy: techniques and applications. Cambridge University Press, 2010. ISBN 978-0-521-86898-3.

[30] Mark D. Wilkinson, Michel Dumontier, IJsbrand Jan Aalbersberg, Gabrielle Appleton, Myles Axton, Arie Baak, Niklas Blomberg, Jan-Willem Boiten, Luiz Bonino da Silva Santos, Philip E. Bourne, Jildau Bouwman, Anthony J. Brookes, Tim Clark, Mercè Crosas, Ingrid Dillo, Olivier Dumon, Scott Edmunds, Chris T. Evelo, Richard Finkers, Alejandra Gonzalez-Beltran, Alasdair J.G. Gray, Paul Groth, Carole Goble, Jeffrey S. Grethe, Jaap Heringa, Peter A.C ‘t Hoen, Rob Hooft, Tobias Kuhn, Ruben Kok, Joost Kok, Scott J. Lusher, Maryann E. Martone, Albert Mons, Abel L. Packer, Bengt Persson, Philippe Rocca-Serra, Marco Roos, Rene van Schaik, Susanna-Assunta Sansone, Erik Schultes, Thierry Sengstag, Ted Slater, George Strawn, Morris A. Swertz, Mark Thompson, Johan van der Lei, Erik van Mulligen, Jan Velterop, Andra Waagmeester, Peter Wittenburg, Katherine Wolstencroft, Jun Zhao, and Barend Mons. The FAIR Guiding Principles for scientific data management and stewardship. Scientific Data, 3(1):160018, March 2016. ISSN 2052-4463. doi:10.1038/sdata.2016.18. URL https://www.nature.com/articles/sdata201618.

[31] M. Okan Irfanoglu, Lindsay Walker, Joelle Sarlls, Stefano Marenco, and Carlo Pierpaoli. Effects of image distortions originating from susceptibility variations and concomitant fields on diffusion MRI tractography results. NeuroImage, 61(1):275–288, May 2012. ISSN 10538119. doi:10.1016/j.neuroimage.2012.02.054. URL https://linkinghub.elsevier.com/retrieve/pii/S1053811912002327.

[32] Céline Provins, Eilidh MacNicol, Saren H. Seeley, Patric Hagmann, and Oscar Esteban. Quality control in functional MRI studies with MRIQC and fMRIPrep. Frontiers in Neuroimaging, 1:1073734, January 2023. ISSN 2813-1193. doi:10.3389/fnimg.2022.1073734. URL https://www.frontiersin.org/articles/10.3389/fnimg.2022.1073734/full.

[33] Richard C. Reynolds, Paul A. Taylor, and Daniel R. Glen. Quality control practices in FMRI analysis: Philosophy, methods and examples using AFNI. Frontiers in Neuroscience, 16:1073800, January 2023. ISSN 1662-453X. doi:10.3389/fnins.2022.1073800. URL https://www.frontiersin.org/articles/10.3389/fnins.2022.1073800/full.

[34] Jerry Allison and Nathan Yanasak. What MRI Sequences Produce the Highest Specific Absorption Rate (SAR), and Is There Something We Should Be Doing to Reduce the SAR During Standard Examinations? American Journal of Roentgenology, 205(2):W140–W140, August 2015. ISSN 0361-803X, 1546-3141. doi:10.2214/AJR.14.14173. URL https://www.ajronline.org/doi/10.2214/AJR.14.14173.

[35] Constantin Sandmann, Erlend Hodneland, and Jan Modersitzki. A practical guideline for T _1_ reconstruction from various flip angles in MRI. Journal of Algorithms & Computational Technology, 10(4):213–223, December 2016. ISSN 1748-3026, 1748-3026. doi:10.1177/1748301816656288. URL http://journals.sagepub.com/doi/10.1177/1748301816656288.

[36] Fabien Balezeau, Pierre-Antoine Eliat, Alejandro Bordelois Cayamo, and Hervé Saint-Jalmes. Mapping of low flip angles in magnetic resonance. Physics in Medicine and Biology, 56(20):6635–6647, October 2011. ISSN 0031-9155, 1361-6560. doi:10.1088/0031-9155/56/20/008. URL https://iopscience.iop.org/article/10.1088/0031-9155/56/20/008.

[37] G Lutterbey, M P Wattjes, J Kandyba, M Harzheim, M V Falkenhausen, N Morakkabati, H Schild, and J Gieseke. Clinical evaluation of a speed optimized T _2_ weighted fast spin echo sequence at 3.0 T using variable flip angle refocusing, half-Fourier acquisition and parallel imaging. The British Journal of Radiology, 80(956):668–673, August 2007. ISSN 0007-1285, 1748-880X. doi:10.1259/bjr/88996134. URL http://www.birpublications.org/doi/10.1259/bjr/88996134.

[38] J. Gonzalez-Castillo, V. Roopchansingh, P.A. Bandettini, and J. Bodurka. Physiological noise effects on the flip angle selection in BOLD fMRI. NeuroImage, 54(4):2764–2778, February 2011. ISSN 10538119. doi:10.1016/j.neuroimage.2010.11.020. URL https://linkinghub.elsevier.com/retrieve/pii/S1053811910014503.

[39] Jingyuan E. Chen and Gary H. Glover. Functional magnetic resonance imaging methods. Neuropsychology Review, 25(3):289–313, Sep 2015. ISSN 1573-6660. doi:10.1007/s11065-015-9294-9.

[40] David A. Feinberg and Kawin Setsompop. Ultra-fast mri of the human brain with simultaneous multi-slice imaging. Journal of Magnetic Resonance, 229:90–100, Apr 2013. ISSN 1090-7807. doi:10.1016/j.jmr.2013.02.002.

[41] Russell A. Poldrack, Paul C. Fletcher, Richard N. Henson, Keith J. Worsley, Matthew Brett, and Thomas E. Nichols. Guidelines for reporting an fmri study. NeuroImage, 40(2):409–414, Apr 2008. ISSN 1053-8119. doi:10.1016/j.neuroimage.2007.11.048.

[42] Samy Ammari, Stephanie Pitre-Champagnat, Laurent Dercle, Emilie Chouzenoux, Salma Moalla, Sylvain Reuze, Hugues Talbot, Tite Mokoyoko, Joya Hadchiti, Sebastien Diffetocq, Andreas Volk, Mickeal El Haik, Sara Lakiss, Corinne Balleyguier, Nathalie Lassau, and Francois Bidault. Influence of Magnetic Field Strength on Magnetic Resonance Imaging Radiomics Features in Brain Imaging, an In Vitro and In Vivo Study. Frontiers in Oncology, 10:541663, January 2021. ISSN 2234-943X. doi:10.3389/fonc.2020.541663. URL https://www.frontiersin.org/articles/10.3389/fonc.2020.541663/full.

[43] Derek K. Jones and Mara Cercignani. Twenty-five pitfalls in the analysis of diffusion MRI data. NMR in Biomedicine, 23(7):803–820, September 2010. ISSN 09523480. doi:10.1002/nbm.1543. URL https://onlinelibrary.wiley.com/doi/10.1002/nbm.1543.

[44] Denis Le Bihan, Cyril Poupon, Alexis Amadon, and Franck Lethimonnier. Artifacts and pitfalls in diffusion MRI. Journal of Magnetic Resonance Imaging, 24(3):478–488, September 2006. ISSN 1053-1807, 1522-2586. doi:10.1002/jmri.20683. URL https://onlinelibrary.wiley.com/doi/10.1002/jmri.20683.

[45] Raúl Tudela, Emma Muñoz-Moreno, Xavier López-Gil, and Guadalupe Soria. Effects of Orientation and Anisometry of Magnetic Resonance Imaging Acquisitions on Diffusion Tensor Imaging and Structural Connectomes. PLOS ONE, 12(1):e0170703, January 2017. ISSN 1932-6203. doi:10.1371/journal.pone.0170703. URL https://dx.plos.org/10.1371/journal.pone.0170703.

[46] Eric Feczko, Greg Conan, Scott Marek, Brenden Tervo-Clemmens, Michaela Cordova, Olivia Doyle, Eric Earl, Anders Perrone, Darrick Sturgeon, Rachel Klein, et al. Adolescent brain cognitive development (abcd) community mri collection and utilities. BioRxiv, pages 2021–07, 2021.

[47] T. Okada, M. Kanagaki, A. Yamamoto, R. Sakamoto, S. Kasahara, E. Morimoto, M. Iima, T. M. Mehemed, S. Nakajima, and K. and Togashi. Which to choose for volumetry: MPRAGE or SPACE? In 2011 ISMRM Annual Meeting and Exhibition (ISMRM), Quebec, Canada, April 2011. ISMRM. URL https://ieeexplore.ieee.org/document/9434081/.

[48] John P. Mugler. Optimized three-dimensional fast-spin-echo mri. ISMRM Annual Meeting and Exhibition, 39(4): 745–767, May 2011. ISSN 1053-1807, 1522-2586. doi:10.1002/jmri.24542.

[49] Patricia Clement, Marco Castellaro, Thomas W. Okell, David L. Thomas, Pieter Vandemaele, Sara Elgayar, Aaron Oliver-Taylor, Thomas Kirk, Joseph G. Woods, Sjoerd B. Vos, Joost P. A. Kuijer, Eric Achten, Matthias J. P. van Osch, John A. Detre, Hanzhang Lu, David C. Alsop, Michael A. Chappell, Luis Hernandez-Garcia, Jan Petr, and Henk J. M. M. Mutsaerts. Asl-bids, the brain imaging data structure extension for arterial spin labeling. Scientific Data, 9(11):543, Sep 2022. ISSN 2052-4463. doi:10.1038/s41597-022-01615-9.

[50] Ben Inglis. A checklist for fMRI acquisition methods reporting in the literature. The Winnower, 2015. ISSN 2373-146X. doi:10.15200/winn.143191.17127. URL https://thewinnower.com/papers/a-checklist-for-fmri-acquisition-methods-reporting-in-the-literature.

[51] Martin A Lindquist and Tor D Wager. Principles of functional Magnetic Resonance Imaging. London: CRC Press, July 2015.

[52] Darcy Mason, Scaramallion, Mrbean-Bremen, Rhaxton Jonathan Suever Vanessasaurus, Dimitri Papadopoulos Orfanos, Guillaume Lemaitre, Aditya Panchal, Alex Rothberg, Markus D. Herrmann, Joan Massich, James Kerns, Korijn Van Golen, Thomas Robitaille, Simon Biggs Moloney, Chris Bridge, Matthew Shun-Shin, Blair Conrad Pawelzajdel, Markus Mattes, YoungKi Lyu, Félix C. Morency, Tim Cogan, Bernardo Pericacho Sánchez, Hans Meine, Joseph Wortmann, Kevin S. Hahn, and Masahiro Wada. pydicom/pydicom: pydicom 2.3.1, November 2022. URL https://zenodo.org/record/7319790.

[53] Pradeep Raamana. Let’s focus our neuroinformatics community efforts in Python and on software validation. https://crossinvalidation.com/2018/05/03/lets-focus-our-neuroinformatics-community-efforts-in-python-and-on-software-validation/, May 2018.

[54] Lisa Scarpace, Adam E. Flanders, Rajan Jain, Tom Mikkelsen, and David W. Andrews. Data From REMBRANDT, 2019. URL https://wiki.cancerimagingarchive.net/display/Public/REMBRANDT. Version Number: 2 Type: dataset.

[55] Lisa Scarpace, Tom Mikkelsen, Soonmee Cha, Sujaya Rao, Sangeeta Tekchandani, David Gutman, Joel H. Saltz, Bradley J. Erickson, Nancy Pedano, Adam E. Flanders, Jill Barnholtz-Sloan, Quinn Ostrom, Daniel Barboriak, and Laura J. Pierce. The Cancer Genome Atlas Glioblastoma Multiforme Collection (TCGA-GBM), 2016. URL https://wiki.cancerimagingarchive.net/x/sgAe. xVersion Number: 4 Type: dataset.

[56] Nancy Pedano, Adam E. Flanders, Lisa Scarpace, Tom Mikkelsen, Jennifer M. Eschbacher, Beth Hermes, Victor Sisneros, Jill Barnholtz-Sloan, and Quinn Ostrom. The Cancer Genome Atlas Low Grade Glioma Collection (TCGA-LGG), 2016. URL https://wiki.cancerimagingarchive.net/x/BANR. xVersion Number: 3 Type: dataset.

[57] Brian K. Rutt and Donald H. Lee. The impact of field strength on image quality in MRI. Journal of Magnetic Resonance Imaging, 6(1):57–62, January 1996. ISSN 10531807, 15222586. doi:10.1002/jmri.1880060111. URL https://onlinelibrary.wiley.com/doi/10.1002/jmri.1880060111.

[58] Alexandre Carré, Guillaume Klausner, Myriam Edjlali, Marvin Lerousseau, Jade Briend-Diop, Roger Sun, Samy Ammari, Sylvain Reuzé, Emilie Alvarez Andres, Théo Estienne, Stéphane Niyoteka, Enzo Battistella, Maria Vakalopoulou, Frédéric Dhermain, Nikos Paragios, Eric Deutsch, Catherine Oppenheim, Johan Pallud, and Charlotte Robert. Standardization of brain MR images across machines and protocols: bridging the gap for MRI-based radiomics. Scientific Reports, 10(1):12340, December 2020. ISSN 2045-2322. doi:10.1038/s41598-020-69298-z. URL http://www.nature.com/articles/s41598-020-69298-z.

[59] Prathyush Chirra, Patrick Leo, Michael Yim, B. Nicolas Bloch, Ardeshir R. Rastinehad, Andrei Purysko, Mark Rosen, Anant Madabhushi, and Satish E. Viswanath. Multisite evaluation of radiomic feature reproducibility and discriminability for identifying peripheral zone prostate tumors on MRI. Journal of Medical Imaging, 6(02):1, June 2019. ISSN 2329-4302. doi:10.1117/1.JMI.6.2.024502. URL https://www.spiedigitallibrary.org/journals/journal-of-medical-imaging/volume-6/issue-02/024502/Multisite-evaluation-of-radiomic-feature-reproducibility-and-discriminability-for-identifying/10.1117/1.JMI.6.2.024502.full.

[60] Marco Bologna, Valentina Corino, and Luca Mainardi. Technical Note: Virtual phantom analyses for preprocessing evaluation and detection of a robust feature set for MRI-radiomics of the brain. Medical Physics, 46(11):5116– 5123, November 2019. ISSN 0094-2405, 2473-4209. doi:10.1002/mp.13834. URL https://onlinelibrary.wiley.com/doi/10.1002/mp.13834.

[61] Niels W. Schurink, Simon R. van Kranen, Sander Roberti, Joost J. M. van Griethuysen, Nino Bogveradze, Francesca Castagnoli, Najim el Khababi, Frans C. H. Bakers, Shira H. de Bie, Gerlof P. T. Bosma, Vincent C. Cappendijk, Remy W. F. Geenen, Peter A. Neijenhuis, Gerald M. Peterson, Cornelis J. Veeken, Roy F. A. Vliegen, Regina G. H. Beets-Tan, and Doenja M. J. Lambregts. Sources of variation in multicenter rectal MRI data and their effect on radiomics feature reproducibility. European Radiology, 32(3):1506–1516, March 2022. ISSN 0938-7994, 1432-1084. doi:10.1007/s00330-021-08251-8. URL https://link.springer.com/10.1007/s00330-021-08251-8.

[62] Yingping Li, Samy Ammari, Corinne Balleyguier, Nathalie Lassau, and Emilie Chouzenoux. Impact of Preprocessing and Harmonization Methods on the Removal of Scanner Effects in Brain MRI Radiomic Features. Cancers, 13(12):3000, June 2021. ISSN 2072-6694. doi:10.3390/cancers13123000. URL https://www.mdpi.com/2072-6694/13/12/3000.

[63] Robert Geirhos, Jörn-Henrik Jacobsen, Claudio Michaelis, Richard Zemel, Wieland Brendel, Matthias Bethge, and Felix A. Wichmann. Shortcut learning in deep neural networks. Nature Machine Intelligence, 2(1111):665–673, Nov 2020. ISSN 2522-5839. doi:10.1038/s42256-020-00257-z.

